# BioTrajX: evaluating biological plausibility in single-cell trajectory inference

**DOI:** 10.64898/2026.11.02.763572

**Authors:** Tsz Ying Li, Yao-Ming Chang, Shih-Yu Chen

## Abstract

**Summary:** Trajectory inference (TI) methods reconstruct cellular progression from single-cell RNA sequencing data, typically by ordering cells along a pseudotime axis. However, different TI methods and analytical choices can produce divergent trajectories from the same dataset, making it difficult to determine which inferred trajectory best reflects the underlying biological process. Existing evaluation strategies often rely on computational benchmarks or qualitative biological assessment, with limited standardized approaches for incorporating *a priori* biological knowledge.

Here, we present BioTrajX, an R package for biologically informed evaluation of pseudotime trajectories using three complementary metrics: Directionality (D), which measures agreement between pseudotime and expected biological progression; Order Consistency (O), which quantifies the coherence of marker-program changes along the trajectory; and Endpoint Validity (E), which assesses enrichment of expected cell states at trajectory endpoints. Together, the DOE metrics provide an interpretable framework for comparing alternative trajectory inference results and evaluating root-cell choices prior to downstream analyses.

**Availability and Implementation:** BioTrajX is implemented in R and freely available under the MIT License at https://github.com/irislity/BioTrajX. Code for reproducing the analyses and figures is available at https://github.com/irislity/BioTrajX_manuscript.

**Supplementary information:** Supplementary data are available at Bioinformatics online.

## 1 INTRODUCTION

Single-cell RNA sequencing (scRNA-seq) enables the characterization of cellular heterogeneity and dynamic biological processes such as development, differentiation, activation, and disease progression. By profiling thousands of individual cells, scRNA-seq captures populations spanning a continuum of molecular states. However, sequencing is destructive and measures different cells at discrete snapshots rather than following the same cells longitudinally. Trajectory inference (TI) methods address this limitation by reconstructing putative biological progression, typically by ordering cells along a pseudotime axis (Saelens *et al*. 2019).

The large and growing number of TI methods makes it challenging for researchers to determine which method is most appropriate for a given biological system. Different algorithms, parameter settings, root selections, and upstream analytical choices can generate substantially different trajectories from the same dataset and alter downstream results (Dong *et al*. 2024). Existing benchmarking frameworks such as dynverse (Saelens *et al*. 2019), scTimeBench (Osakwe, Huang, and Li 2026), and Escort (Dong *et al*. 2024) evaluate trajectory or temporal modelling methods using computational, reference-based, and data-driven criteria, with limited incorporation of *a priori* biological knowledge. Agreement with such benchmarks does not necessarily establish whether an inferred trajectory is consistent with the expected biological progression of the system under study. Because scRNA-seq provides destructively sampled snapshots rather than direct observations of cellular transitions, definitive trajectory ground truth is often unavailable. Researchers may therefore rely heavily on prior biological knowledge and subjective visual assessment to judge whether an inferred trajectory is biologically plausible and to select among alternative trajectories (Dong *et al*. 2024).

A fundamental assumption of TI is that inferred pseudotemporal ordering should reflect underlying biological progression. Trajectory quality is therefore commonly assessed by examining whether known marker programs change in the expected direction and whether annotated cellular states occupy biologically plausible positions along the trajectory. Indeed, widely used TI methods including Monocle (Trapnell *et al*. 2014), diffusion pseudotime (DPT) (Haghverdi *et al*. 2016), Palantir (Setty *et al*. 2019), and Slingshot (Street *et al*. 2018) were originally validated, in part, by recovering expected marker dynamics or positioning known cell populations along biologically meaningful trajectories.

However, these biological assessments are generally qualitative and lack a standardized quantitative framework for comparing alternative trajectories. To address this gap, we developed BioTrajX, an R package for biologically informed trajectory evaluation using three complementary metrics: Directionality (D), which assesses expected marker-program direction; Order Consistency (O), which quantifies coherent marker progression; and Endpoint Validity (E), which evaluates recovery of expected endpoint states (Fig. 1A). BioTrajX operates independently of the TI algorithm and enables researchers to compare alternative methods and root-cell choices before downstream biological analyses.

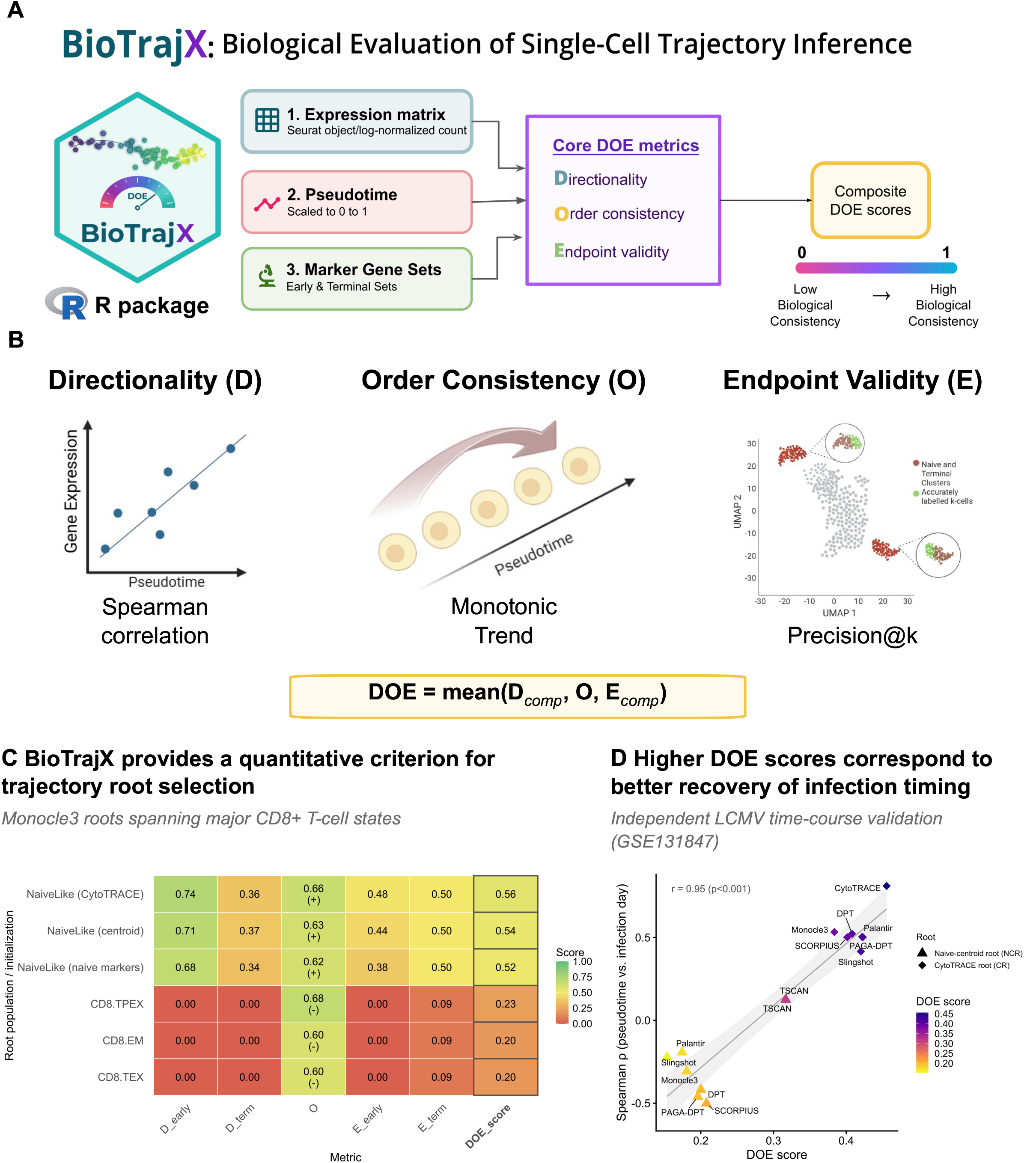

## 2 IMPLEMENTATION

BioTrajX provides a unified framework for biologically evaluating trajectory inference results using three complementary metrics: Directionality (D), Order Consistency (O), and Endpoint Validity (E). It integrates with Seurat, supports user-defined and curated marker sets, and provides interpretable diagnostics for trajectory orientation, marker ordering, and endpoint recovery (Fig. 1B). The package supports both linear and branched trajectories, enabling systematic evaluation of continuous developmental processes and lineage bifurcations. The workflow consists of three steps: (i) loading the required inputs, (ii) computing the DOE metrics, and (iii) visualizing and comparing trajectory inference results.

### 2.1 Input and Workflow

BioTrajX accepts three required inputs: a log-normalized expression matrix, pseudotime values, and predefined early and terminal marker gene sets. Pseudotime values are internally rescaled to the ([0,1]) interval using min–max normalization before metric calculation. Expression data can be supplied as a Seurat object or any log-normalized dense (matrix) or sparse (dgCMatrix) matrix. Marker sets may be user-defined or obtained from curated resources such as MSigDB (Subramanian et al. 2005) or CellMarker 2.0 (Hu *et al*. 2023), allowing evaluation across diverse biological systems. Optional cell-state labels or cluster annotations can be provided for endpoint assessment.

For branched trajectories, branch membership is specified using user-defined cell type or cluster annotations, enabling branch-specific evaluation while preserving a shared pseudotime ordering.

### 2.2 DOE Metrics

BioTrajX quantifies complementary aspects of trajectory quality using three biologically informed metrics. Directionality (D) evaluates whether early and terminal marker programs change in the expected biological direction along pseudotime using Spearman correlation. The Spearman correlation coefficient ρ captures the coarse, global trend between marker expression and pseudotime, indicating whether a marker program generally increases or decreases along the trajectory. Order Consistency (O) complements this global measure by quantifying finer-grained monotonic progression across pseudotime using direction-constrained isotonic regression applied to pseudobulk expression. The isotonic regression R^2^ measures how consistently successive pseudotime bins conform to the expected monotonic trend and is calibrated against a simulated null distribution. Scores near 1 indicate strong monotonic ordering, whereas scores near 0 indicate flat or random expression patterns. An accompanying orientation sign (+/−) distinguishes correctly oriented from reversed trajectories. Endpoint Validity (E) assesses enrichment of expected early and terminal cell states at trajectory endpoints using ranked-cell precision. Endpoint states can be specified using user-provided labels or inferred automatically from marker score distributions using two-component Gaussian mixture modelling. A composite DOE score is calculated as the arithmetic mean of D, O, and E, providing an overall summary of biological trajectory quality.

### 2.3 Outputs and Visualization

BioTrajX returns the individual D, O, and E metrics together with a composite DOE score for each inferred trajectory. For branched trajectories, metrics are computed independently for each lineage and summarized across branches.

Built-in visualization functions generate comparative heatmaps, bar charts, radar plots, and metric-specific diagnostic plots, facilitating systematic comparison of trajectory inference results while providing interpretable diagnostics for trajectory orientation, marker ordering, and endpoint recovery.

## 3 APPLICATIONS

### 3.1 Controlled Validation Using Simulated Datasets

We first evaluated BioTrajX on synthetic datasets with known marker programs to validate metric behaviour under controlled conditions. Zero-inflated and weak-signal datasets produced intermediate scores, whereas flat, high-dropout, and shuffled-pseudotime controls were strongly penalized (Fig. S1). Notably, shuffling pseudotime reduced the DOE score of the ZINB trajectory from 0.81 to 0.13 in the linear setting and from 0.79 to 0.12 in the branched setting, despite preserving the underlying expression distributions, indicating that DOE effectively rewards trajectories with better agreement between marker dynamics and correct pseudotime ordering.

These synthetic datasets collectively illustrate that the three BioTrajX components capture complementary failure modes (Fig. S2-S7). Order consistency can remain high when a trajectory is globally reversed, whereas Directionality and Endpoint validity distinguish whether the inferred ordering is biologically oriented and recovers the expected terminal states. Conversely, disruption of the ordering itself through pseudotime shuffling causes all three components to approach null-like values. Thus, a high score in any single component is insufficient to establish biological plausibility. The composite DOE score integrates complementary evidence about trajectory orientation, marker ordering, and endpoint recovery.

### 3.2 Benchmarking Trajectory Inference Results Across Real Datasets

We next asked whether BioTrajX could distinguish biologically plausible trajectory solutions produced by eight commonly used TI methods: Slingshot (Street et al. 2018), CytoTRACE (Gulati et al. 2020), Monocle3 (Cao et al. 2019), DPT (Haghverdi et al. 2016), SCORPIUS (Cannoodt et al. 2016), PAGA-DPT (Wolf et al. 2019), Palantir (Setty et al. 2019), and TSCAN (Ji and Ji 2016). We evaluated the relationship between DOE score and an independent temporal ground-truth reference before applying BioTrajX to datasets representing linear and branched biological trajectories.

It was previously shown that Slingshot inferred a reversed differentiation trajectory in a longitudinal LCMV infection dataset with known sampling times (Hudson and Wieland, 2023). Using the same dataset (GSE131847), we asked whether BioTrajX could distinguish trajectories that recover the experimentally observed progression and whether its evaluation remained informative across alternative root choices (Fig. S8). Trajectory configurations were evaluated using either the CytoTRACE-selected root (CR) or a naive-centroid root (NCR). Under the CR configuration, the inferred trajectories showed positive orientation signs, whereas most NCR configurations showed the opposite orientation; TSCAN retained a positive orientation under both configurations (Fig. S8C). CytoTRACE achieved the highest DOE score and the strongest positive association between pseudotime and day post-infection, while lower-scoring configurations generally showed weaker or reversed temporal relationships (Fig. 1C; Fig. S8D). Across method and root configurations, DOE score was strongly associated with temporal recovery (Pearson’s r=0.95, p<0.001; Fig. 1C), quantified independently as the Spearman correlation between pseudotime and experimental day. These results show that BioTrajX can distinguish biologically consistent from incorrectly oriented trajectory solutions across both method and root-selection choices without using experimental sampling time as an evaluation input, while its component metrics further indicate whether the discrepancy arises from reversed ordering, inconsistent marker directionality, or misplaced trajectory endpoints, thereby helping diagnose root-dependent failures in pseudotime inference.

We next evaluated a linear CD8+ T-cell exhaustion dataset (Kürten et al. 2021), in which cells are expected to progress broadly from naive-like states through effector populations toward progenitor and terminal exhaustion (Fig. S9). BioTrajX revealed substantial variation in the extent to which TI methods reproduced this progression. TSCAN achieved the highest composite DOE score (0.60), followed by Monocle3 (0.54), PAGA-DPT (0.52), CytoTRACE and Slingshot (0.49), Palantir (0.48), SCORPIUS (0.46), and DPT (0.41) (Fig. S9C). Examination of marker-score trends, exhaustion-associated transcription factors, and pseudotime distributions across annotated cell states showed that trajectories with higher DOE scores more clearly recapitulated the expected naive-to-exhausted transition (Fig. S9B, D, E).

The branched haematopoietic differentiation dataset provided a complementary test because a single global pseudotime cannot adequately describe both lineage outcomes. We therefore evaluated Stem-to-Erythroid and Stem-to-B trajectories independently (Fig. S10). In the erythroid branch, CytoTRACE showed the highest biological consistency (DOE = 0.80), followed by DPT (0.71), Palantir (0.64), and SCORPIUS (0.63). In contrast, the B-cell branch was ranked highest for Slingshot (0.59), followed by PAGA-DPT (0.57), Palantir (0.53), and DPT (0.53). Marker-score trajectories, lineage-associated TF expression, and pseudotime distributions across developmental stages showed corresponding differences in how well each inferred trajectory recapitulated the expected progression within each branch (Fig. S10D–H). The differing DOE rankings between the erythroid and B-cell branches further showed that the biological plausibility of a given TI solution can vary across lineage contexts. These analyses illustrate how BioTrajX can be used to compare alternative trajectory solutions at the branch level, allowing distinct lineage outcomes to be evaluated against their own expected biological transitions.

### 3.3 Quantitative Evaluation of Root Cell Selection

Root specification represents a common source of analyst-dependent variation in trajectory inference. Most TI methods require a starting population or root to orient pseudotime. In practice, roots are frequently chosen using prior biological knowledge or visual inspection, introducing a subjective step into the analysis.

We therefore asked whether BioTrajX could provide a quantitative criterion for comparing candidate roots. Using Monocle3 on the CD8+ T-cell exhaustion dataset, we evaluated roots spanning the major annotated cell states (Fig. 1D; Fig. S11). All three roots selected from the CD8.NaiveLike population produced substantially higher DOE scores, including the CytoTRACE-selected cell (DOE = 0.56), the cluster-centroid cell (0.54), and the cell with the highest naive-marker score (0.52). In contrast, roots placed in later differentiation states produced markedly lower scores, with CD8.TPEX, CD8.EM, and CD8.TEX roots yielding DOE scores of 0.23, 0.20, and 0.20, respectively. This analysis illustrates a second use of BioTrajX as a quantitative tool for comparing alternative trajectory initializations, with roots placed in the expected naive-like starting population consistently producing higher DOE scores than roots placed in later differentiation states.

## 4 CONCLUSION

BioTrajX complements existing computational benchmarking frameworks by incorporating biological prior knowledge into a quantitative framework for trajectory evaluation. Through three complementary metrics: Directionality (D), Order Consistency (O), and Endpoint Validity (E), it enables systematic evaluation of trajectory inference results while providing interpretable diagnostics for biological assessment, supporting objective comparison and selection of inferred trajectories and root cell selection prior to downstream analyses such as differential expression, gene regulatory network inference, lineage analysis, and benchmarking of new trajectory inference methods.

## 5 ACKNOWLEDGEMENTS

We thank the Academia Sinica Grid-computing Center (ASGC) for providing Distributed Cloud computing resources supported by Academia Sinica. We also thank Hung-Hsin Chen for suggesting the use of simulated datasets to validate the BioTrajX metrics.

