## supplementary_material for "BioTrajX: evaluating biological plausibility in single-cell trajectory inference"

### Supplementary Materials for "BioTrajX: evaluating biological plausibility in single-cell trajectory inference"

**Abstract**

This supplementary document provides detailed definitions of the DOE metrics, benchmark dataset construction procedures, and additional validation results supporting the BioTrajX framework.

### Supplementary Methods S1. Mathematical definition of DOE metrics

Let $\mathcal{C}$ denote the set of cells and let $\mathbf{t} = (t_i)_{i\in\mathcal{C}}$ denote the pseudotime assignment, scaled to the interval $[0, 1]$. Let $\mathcal{G}_N$ and $\mathcal{G}_T$ denote the early and terminal marker-gene sets used for evaluation, respectively. For a marker set $\mathcal{F}$, BioTrajX computes a per-cell marker score as

$$S_i^{(\mathcal{F})} = \frac{1}{|\mathcal{F}|} \sum_{f\in\mathcal{F}} z_{if}, \tag{1}$$

where $z_{if}$ is the standardized expression of marker gene $f$ in cell $i$. These marker scores are used to evaluate whether the inferred pseudotime is consistent with expected biological progression.

### Directionality

Directionality provides a coarse-grained, program-level assessment of trajectory orientation. It evaluates whether early and terminal marker programs change in the expected direction along pseudotime. Because marker dynamics may be nonlinear or saturated, BioTrajX uses Spearman rank correlation without assuming a linear relationship between marker score and pseudotime.

Early marker scores are expected to decrease, whereas terminal marker scores are expected to increase:

$$D_N = \max\Big(0,\ -\rho_S\Big(\mathbf{t}, S^{(N)}\Big)\Big), \tag{2}$$

$$D_T = \max\Big(0,\ \ \rho_S\Big(\mathbf{t}, S^{(T)}\Big)\Big), \tag{3}$$

where $\rho_S$ denotes Spearman correlation, $S^{(N)}$ is the per-cell early marker score, and $S^{(T)}$ is the per-cell terminal marker score. The opposing signs reflect the expected decline of early marker scores and rise of terminal marker scores along pseudotime. Both $D_N$ and $D_T$ are bounded between 0 and 1, with higher values indicating stronger agreement with the expected direction of biological progression.

Thus, Directionality captures the global orientation of the trajectory at the marker-program level, rather than requiring every individual marker gene to follow the expected pattern.

### Order Consistency

Order Consistency provides a finer-grained, gene-level assessment of biological ordering. Whereas Directionality evaluates the overall orientation of aggregate marker programs, Order Consistency evaluates each marker gene individually, quantifying whether its expression follows the expected monotonic progression along pseudotime. Each marker gene $g$ is assigned an expected direction,

$$\Delta_g = \begin{cases} -1, & g \in \mathcal{G}_N, \\ +1, & g \in \mathcal{G}_T, \end{cases}$$

where $\mathcal{G}_N$ and $\mathcal{G}_T$ denote early/root and terminal marker genes, respectively.

To reduce cell-level noise and dropout, cells are grouped into $K$ equal-width pseudotime bins. For each marker gene, BioTrajX computes the pseudobulk mean expression $\bar{x}_{gk}$ in each bin. Marker profiles are then oriented according to their expected direction,

$$\tilde{x}_{gk} = \Delta_g \bar{x}_{gk},$$

so that biologically consistent trends are represented as non-decreasing profiles.

For each retained marker gene, BioTrajX fits a non-decreasing isotonic regression curve to the oriented binned expression profile:

$$\hat{\mathbf{x}}_g = \arg\min_{\mathbf{y}\in\mathcal{I}} \sum_{k=1}^{K} (\tilde{x}_{gk} - y_k)^2,$$

where $\mathcal{I}$ is the set of non-decreasing sequences over ordered pseudotime bins. The raw monotonic fit is measured by

$$R_g^2 = 1 - \frac{\sum_{k=1}^{K} (\tilde{x}_{gk} - \hat{x}_{gk})^2}{\sum_{k=1}^{K} (\tilde{x}_{gk} - \bar{\tilde{x}}_g)^2}.$$

Genes with no variation across bins are assigned a score of zero.

Because isotonic regression can produce non-zero fits even for random profiles, BioTrajX calibrates the raw score against a simulated null expectation, $R^2_{\mathrm{null}}(K)$. The calibrated per-gene Order Consistency score is

$$O_g = \max\left(0, \frac{R_g^2 - R^2_{\mathrm{null}}(K)}{1 - R^2_{\mathrm{null}}(K)}\right).$$

The final Order Consistency score is the mean across retained marker genes:

$$O = \frac{1}{|\mathcal{G}^*|} \sum_{g \in \mathcal{G}^*} O_g,$$

where $\mathcal{G}^*$ denotes marker genes retained after variability filtering. By default, BioTrajX evaluates both pseudotime orientations and reports the orientation with the higher Order Consistency score. Higher values indicate stronger gene-resolved consistency between marker expression and the expected monotonic progression along pseudotime, thereby capturing finer-scale ordering that may be obscured by aggregate marker-program scores.

### Endpoint Validity

Endpoint Validity evaluates whether the extremes of pseudotime correspond to expected biological endpoint states. Let $\mathcal{R}$ and $\mathcal{T}$ denote cells labeled as early/root and terminal, respectively, with $k_N = |\mathcal{R}|$ and $k_T = |\mathcal{T}|$.

For the early endpoint, BioTrajX selects the first $k_N$ cells ranked by ascending pseudotime and computes

$$E_N = \frac{|\{i \in \mathcal{R} : \mathrm{rank}_{\mathrm{asc}}(t_i) \leq k_N\}|}{k_N}.$$

For the terminal endpoint, BioTrajX selects the first $k_T$ cells ranked by descending pseudotime and computes

$$E_T = \frac{|\{i \in \mathcal{T} : \mathrm{rank}_{\mathrm{desc}}(t_i) \leq k_T\}|}{k_T}.$$

Thus, $E_N$ and $E_T$ correspond to precision at $k_N$ and $k_T$, respectively, and range from 0 to 1.

Endpoint labels can be supplied from cluster annotations or inferred from marker scores. In marker-score mode, BioTrajX fits a two-component Gaussian mixture model to each endpoint marker-score distribution,

$$p(s_i) = \sum_{m=1}^{2} \pi_m \mathcal{N}(s_i \mid \mu_m, \sigma_m^2),$$

and labels cells assigned to the higher-mean component as candidate endpoint cells. If mixture fitting is unstable or produces invalid labels, BioTrajX uses fallback strategies, including quantile-based splitting, two-means clustering, or thresholding at the empirical mean plus one half standard deviation. The labeling method is recorded in the output for diagnostic interpretation.

#### Composite DOE Score

BioTrajX computes a composite DOE score as the arithmetic mean of the available component scores, providing a single summary measure of overall trajectory quality on the interval $[0,1]$.

D, O, E capture conceptually distinct properties of an inferred trajectory and are therefore weighted equally in the composite score:

$$\text{DOE score} = \frac{D_{\text{comp}} + O + E_{\text{comp}}}{3}, \quad \text{where} \quad D_{\text{comp}} = \frac{2}{\frac{1}{D_N} + \frac{1}{D_T}} \quad \text{and} \quad E_{\text{comp}} = \frac{2}{\frac{1}{E_N} + \frac{1}{E_T}}.$$

O is orientation-invariant, reversed trajectories may still receive high scores. Therefore, the orientation sign should therefore be inspected alongside O to distinguish correctly oriented trajectories from strongly ordered but reversed trajectories.

### Supplementary Methods S2. Simulated Datasets

#### Synthetic Datasets

Synthetic expression matrices were generated to evaluate the expected behavior of the DOE metrics under controlled trajectory scenarios. All datasets used $G = 50$ genes (or $G = 40$ for branched datasets) and $N = 1{,}000$ cells per linear scenario or $N = 500$ cells per branch (1,500 total) for branched scenarios. The simulation framework included nine linear and six branched scenarios. Linear datasets evaluated performance across idealized monotonic trajectories, reversed orientations, abrupt state transitions, realistic single-cell count distributions, and challenging conditions characterised by weak biological signal or extensive dropout. Branched datasets extended these scenarios to an $A \to B$ and $A \to C$ bifurcation topology.

Table 1 summarises the nine linear datasets, and Table 2 summarises the six branched datasets.

Table 1: Linear synthetic datasets ($G = 50$, $N = 1{,}000$).

| Dataset | Noise model | $\mu_{\text{early}}(t)$ | $\mu_{\text{term}}(t)$ | Purpose |
|---|---|---|---|---|
| Gaussian linear | Gaussian ($\sigma = 0.3$) | $8 - 7t$ | $1 + 7t$ | Performance ceiling / positive control |
| Gaussian reversed | Gaussian ($\sigma = 0.3$) | (same expression, pseudotime reversed) | - | Negative control: pseudotime reversed |
| Stepwise repression | ZINB ($\theta = 1,\ \pi_0 = 0.2$) | $\begin{cases} 8, & t \le 0.25 \\ 1, & \text{otherwise} \end{cases}$ | $\begin{cases} 8, & t \ge 0.75 \\ 1, & \text{otherwise} \end{cases}$ | Abrupt endpoint marker programs |
| Stepwise activation | ZINB ($\theta = 1,\ \pi_0 = 0.2$) | (same expression, pseudotime reversed) | - | Reversed endpoint orientation |
| ZINB linear | ZINB ($\theta = 1.5,\ \pi_0 = 0.05$) | $8 - 7t$ | $1 + 7t$ | Baseline scRNA-seq simulation |
| ZINB reversed | ZINB ($\theta = 1.5,\ \pi_0 = 0.05$) | $8 - 7t$ | $1 + 7t$ | Orientation robustness test |
| Flat / no signal | ZINB ($\theta = 0.5,\ \pi_0 = 0.05$) | 2 | 2 | Uninformative trajectory baseline |
| High dropout | ZINB ($\theta = 0.5,\ \pi_0 = 0.9$) | $8 - 7t$ | $1 + 7t$ | Robustness to extreme sparsity |
| Low signal | ZINB ($\theta = 0.5,\ \pi_0 = 0.5$) | $1.5 - 0.5t$ | $1 + 0.5t$ | Sensitivity near the noise floor |

Table 2: Branched synthetic datasets ($A \to B$ and $A \to C$ topology; $G = 40$, $N = 500$ cells per branch, 1,500 total). Early markers (genes 1–10) are shared; terminal markers (genes 11–20 for branch $B$; genes 21–30 for branch $C$).

| Dataset | Noise model | $\mu_{\text{early}}(t)$ | $\mu_{\text{term}}(t)$ | Purpose |
|---|---|---|---|---|
| Gaussian branched | Gaussian ($\sigma = 0.3$) | $8 - 7t$ | $1 + 7t$ (within branch) | Branched ceiling / positive control |
| Gaussian branched reversed | Gaussian ($\sigma = 0.3$) | (same expression, pseudotime reversed) | - | Negative control: branched reversed |
| Branched stepwise repression | ZINB ($\theta = 1,\ \pi_0 = 0.2$) | $\begin{cases} 8, & t \le 0.25 \\ 1, & \text{otherwise} \end{cases}$ | Branch-specific step at $t \ge 0.75$ | Abrupt endpoint programs, branched |
| Branched stepwise activation | ZINB ($\theta = 1,\ \pi_0 = 0.2$) | (same expression, pseudotime reversed) | - | Reversed endpoint orientation, branched |
| Branched ZINB | ZINB ($\theta = 1.5,\ \pi_0 = 0.05$) | $8 - 6t$ | $2 + 6u$ (within branch) | Realistic scRNA-seq, branched |
| Branched low signal | ZINB ($\theta = 0.5,\ \pi_0 = 0.5$) | $1.5 - 0.5t$ | $1 + 0.5u$ (within branch) | Sensitivity near noise floor, branched |

Here $t \in [0,1]$ is global pseudotime and $u \in [0,1]$ is pseudotime rescaled within the terminal branch ($u = (t - 0.5)/0.5$ for $t \in [0.5, 1]$).

**Gaussian Linear** The Gaussian linear dataset creates an ideal early-to-late trajectory with $G = 50$ genes and $N = 1{,}000$ cells. Cells are assigned pseudotime $t_i \in [0,1]$. Genes 1–10 are early markers that decrease along pseudotime, genes 11–20 are terminal markers that increase along pseudotime, and genes 21–50 are constant background genes.

The expected expression profile for each gene is defined as

$$\mu_{gi} = \begin{cases} (1+s) - st_i, & g = 1, \dots, 10 \quad \text{(early markers)}, \\ 1 + st_i, & g = 11, \dots, 20 \quad \text{(terminal markers)}, \\ 2, & g = 21, \dots, 50 \quad \text{(background genes)}. \end{cases}$$

Observed expression values are generated by adding Gaussian noise

$$X_{gi} = \mu_{gi} + \epsilon_{gi}, \qquad \epsilon_{gi} \sim \mathcal{N}(0, \sigma^2),$$

followed by truncation of negative values to zero. We used $s = 7$ and $\sigma = 0.3$, yielding marker expression ranges from 1 to 8, with noise small relative to the dynamic range of the marker signal.

**Gaussian Reversed** The Gaussian reversed dataset was generated from the Gaussian linear dataset by reversing the assigned pseudotime values while keeping the expression matrix unchanged:

$$\tilde{t}_i = 1 - t_i.$$

This dataset tests whether BioTrajX correctly penalizes trajectories with the opposite orientation relative to the known early-to-terminal marker program.

**Zero-Inflated Negative Binomial Sampling** For ZINB-based datasets, raw counts were sampled from a zero-inflated negative binomial distribution:

$$C_{gi} \sim \text{ZINB}(\mu_{gi}, \theta, \pi_0),$$

where $\mu_{gi}$ is the mean expression, $\theta$ is the negative binomial dispersion parameter, and $\pi_0$ is the zero-inflation probability. Equivalently,

$$C_{gi} = \begin{cases} 0, & \text{with probability } \pi_0, \\ X_{gi}, & \text{with probability } 1 - \pi_0, \end{cases} \qquad X_{gi} \sim \text{NB}(\mu_{gi}, \theta).$$

The final expression matrix contains log-transformed values:

$$E_{gi} = \log(1 + C_{gi}).$$

**Stepwise Repression** The stepwise repression dataset creates a trajectory in which early-state marker expression is concentrated at the start of pseudotime and terminal-state expression is concentrated at the end, mimicking abrupt cell-state transitions. We used $N = 1{,}000$ cells, $G = 50$ genes, $q_n = q_t = 0.25$, $h = 8$, $\ell = 1$, $\theta = 1$, and $\pi_0 = 0.2$. The expected expression means are

$$\mu_{gi} = \begin{cases} h \text{ if } t_i \le q_n, \ \ell \text{ otherwise}, & g = 1, \dots, 10, \\ h \text{ if } t_i \ge 1 - q_t, \ \ell \text{ otherwise}, & g = 11, \dots, 20, \\ 2, & g = 21, \dots, 50. \end{cases}$$

Observed expression values are sampled from $X_{gi} \sim \text{ZINB}(\mu_{gi}, \theta, \pi_0)$.

**Stepwise Activation** The stepwise activation dataset uses the same simulated expression matrix as the stepwise repression dataset, but with the pseudotime vector reversed:

$$t_i^{\text{act}} = 1 - t_i.$$

As a result, true terminal-state cells appear at $t = 0$ and true early-state cells appear at $t = 1$, so all endpoint and directionality scores are expected to be near zero.

**ZINB Linear** The ZINB linear dataset uses the same monotonic mean structure as the Gaussian linear dataset, but with ZINB noise ($\theta = 1.5$, $\pi_0 = 0.05$) to mimic baseline scRNA-seq count distributions:

$$\mu_{gi} = \begin{cases} 8 - 7t_i, & g = 1, \dots, 10, \\ 1 + 7t_i, & g = 11, \dots, 20, \\ 2, & g = 21, \dots, 50. \end{cases}$$

**ZINB Reversed** The ZINB reversed dataset uses the same expression matrix as the ZINB linear dataset, but the pseudotime vector is reversed:

$$t_i^{\text{rev}} = 1 - t_i.$$

**Flat / No Signal** The flat dataset creates an uninformative trajectory in which all genes are sampled from the same expression distribution, independent of pseudotime:

$$\mu_{gi} = 2, \qquad g = 1, \dots, G, \quad i = 1, \dots, N.$$

We used $\theta = 0.5$ and $\pi_0 = 0.05$. Genes 1–10 and 11–20 were still designated as early and terminal marker sets, respectively, but no trajectory-correlated signal was present.

**High-Dropout and Low-Signal Datasets** The high-dropout and low-signal datasets are stress-test variants of the ZINB linear dataset. Both used the same monotonic mean structure:

$$\mu_{gi} = \begin{cases} (1 + s) - st_i, & g = 1, \dots, 10, \\ 1 + st_i, & g = 11, \dots, 20, \\ 2, & g = 21, \dots, 50. \end{cases}$$

The high-dropout dataset used $s = 7$, $\theta = 0.5$, and $\pi_0 = 0.9$ to assess robustness under extreme sparsity (approximately 90% zeros). The low-signal dataset used $s = 0.5$, $\theta = 0.5$, and $\pi_0 = 0.5$ to assess metric sensitivity when the marker gradient is weak (mean range 1 to 1.5).

**ZINB linear, shuffled** Uses the same expression matrix as the ZINB linear dataset ($\theta = 1.5$, $\pi_0 = 0.05$, $N = 1{,}000$), with pseudotime independently shuffled across all cells. Expected: $D$, $O$, and $E$ all collapse toward the noise floor ($\approx 0$).

#### Branched Datasets

Branched datasets used an $A \to B$ and $A \to C$ bifurcation topology with $G = 40$ genes and $N = 500$ cells per branch (1,500 total). Cells in branch $A$ were assigned global pseudotime $t \in [0, 0.5]$; cells in branches $B$ and $C$ were assigned $t \in [0.5, 1.0]$. Genes 1–10 were shared early markers (decreasing along global pseudotime in all cells); genes 11–20 were $B$-specific terminal markers (increasing only within branch $B$); genes 21–30 were $C$-specific terminal markers (increasing only within branch $C$); genes 31–40 were background genes with constant mean expression. DOE was computed separately for the $A \to B$ and $A \to C$ trajectories.

**Gaussian branched** Used the same linear mean structure as the Gaussian linear dataset ($s = 7$, $\sigma = 0.3$), providing a high-SNR ceiling reference for the branched setting. Early markers followed $\mu = 8 - 7t$ across all cells; branch-specific terminal markers followed $\mu = 1 + 7t$ within their respective branches using global pseudotime.

**Gaussian branched reversed** Same expression matrix as the Gaussian branched dataset with pseudotime reversed ($\tilde{t}_i = 1 - t_i$). Expected per-branch: $D \approx 0$, $E \approx 0$.

**Branched stepwise repression** Used the same step-function mean structure as the linear stepwise repression dataset ($q_n = q_t = 0.25$, $h = 8$, $\ell = 1$, $\theta = 1$, $\pi_0 = 0.2$), with branch-specific terminal markers expressed only in the final 25% of pseudotime within each terminal branch. Expected per-branch: $E_N \approx 1$, $E_T \approx 1$.

**Branched stepwise activation** Same expression matrix as the branched stepwise repression dataset with pseudotime reversed. True endpoints appear at the wrong ends of pseudotime. Expected per-branch: $E_N \approx 0$, $E_T \approx 0$.

**Branched ZINB** Used ZINB noise ($\theta = 1.5$, $\pi_0 = 0.05$) matching the ZINB linear dataset. Early markers followed $\mu = 8 - 6t$ across all cells; branch-specific terminal markers followed $\mu = 2 + 6u$ within their respective branches, where $u = (t - 0.5)/0.5 \in [0, 1]$ is pseudotime rescaled within the terminal branch.

**Branched low signal** Used ZINB noise ($\theta = 0.5$, $\pi_0 = 0.5$) and a weak gradient ($s = 0.5$). Early markers followed $\mu = 1.5 - 0.5t$; branch-specific terminal markers followed $\mu = 1 + 0.5u$. This dataset tests metric sensitivity near the noise floor in branched trajectories.

**Branched ZINB, shuffled** Uses the same expression matrix as the branched ZINB dataset ($\theta = 1.5$, $\pi_0 = 0.05$), with pseudotime independently shuffled across all $N = 1{,}500$ cells pooled across branches $A$, $B$, and $C$, destroying both the within-branch ordering and the $A \to B$ / $A \to C$ branch structure. Expected per-branch: $D$, $O$, and $E$ all collapse toward the noise floor ($\approx 0$).

### Supplementary Methods S3. Real Datasets

This section describes the real single-cell datasets used to assess BioTrajX on experimentally observed biological trajectories.

##### Trajectory Inference Wrapper

To benchmark trajectory inference (TI) methods on real single-cell datasets, we implemented a unified wrapper for eight methods: Slingshot, CytoTRACE, Monocle3, DPT, SCORPIUS, TSCAN, PAGA–DPT, and Palantir. Each method produced a single cell-level pseudotime vector for downstream BioTrajX evaluation. The primary expression input comprised the top 2,000 highly variable genes from the normalised expression matrix; an all-gene matrix was supplied separately to CytoTRACE to retain information on gene detection across cells. Precomputed Seurat PCA embeddings were reused by Slingshot, DPT, PAGA–DPT, and Palantir. Slingshot used at most the first 10 principal components, whereas PAGA–DPT and Palantir used at most the first 30; DPT used the supplied embedding. An R random seed of 42 was set before the joint TI run.

**Root cell selection** For each dataset, the trajectory root was selected from a biologically defined early population (e.g. naive cells, `CD8.NaiveLike`, or `Stem_Progenitors`). Using the first $d = \min(20, D)$ principal components, we calculated the centroid of this population,

$$\boldsymbol{\mu} = \frac{1}{|\mathcal{N}|} \sum_{i \in \mathcal{N}} \mathbf{x}_i, \tag{4}$$

and selected the observed cell nearest to the centroid,

$$c^* = \arg\min_{i \in \mathcal{N}} \|\mathbf{x}_i - \boldsymbol{\mu}\|_2 \,. \tag{5}$$

This nearest-centroid root (NCR) provides a representative, method-agnostic cell from the predefined early population. The resulting cell $c^*$ was supplied as the fixed `start_cell` argument to all root-dependent trajectory-inference methods (Slingshot, Monocle 3, DPT, SCORPIUS, TSCAN, PAGA-DPT, and Palantir). For SCORPIUS and TSCAN, trajectory orientation was aligned with the selected root cell after pseudotime rescaling. If the root had an available pseudotime greater than 0.5, all pseudotime values were replaced by $1 - t_i$, placing the root in the earlier half of the pseudotime range.

CytoTRACE itself did not require an externally specified root. Where a CytoTRACE-based root (CR) was evaluated as a sensitivity configuration, the candidate was selected from the same predefined early population as the NCR. CytoTRACE scores were inverted and rescaled so that lower values represented greater developmental potential, and the cell with the lowest resulting value was selected as the CR.

For Slingshot, cells were partitioned by $k$-means clustering in PCA space, with $k = \min\left(10, \max\left(3, \mathrm{round}\left(\frac{\sqrt{n}}{3}\right)\right)\right)$, where $n$ denotes the number of cells. Clustering used 25 random starts and a maximum of 50 iterations. Slingshot then inferred lineages and fitted principal curves, starting from the cluster containing the selected root cell. Lineage-specific pseudotimes were collapsed to a single value per cell using a weighted mean of the available pseudotimes, with weights obtained from the fitted curves. Cells with zero total lineage weight were assigned missing values.

DPT was computed using the `destiny` package by constructing a diffusion map from the PCA embedding and specifying the selected root cell as the starting tip. Monocle3 performed preprocessing with up to 30 dimensions, generated a UMAP embedding, clustered cells, and learned a principal graph with `use_partition = FALSE`. Cells were then ordered from the selected root, and infinite pseudotimes were converted to missing values. SCORPIUS performed dimensionality reduction using Spearman dissimilarities and inferred a one-dimensional trajectory. TSCAN performed expression-based clustering with candidate cluster numbers from 2 to 9 and obtained a cell ordering using `TSCANorder`. Positions in the returned ordering were converted to pseudotime ranks; cells absent from that ordering were assigned missing values.

PAGA–DPT and Palantir were executed in isolated Python subprocesses to avoid conflicts between R and Python numerical libraries. For PAGA–DPT, Scanpy constructed a nearest-neighbour graph with

15 neighbours from the PCA embedding, performed Leiden clustering at resolution 0.5, and computed PAGA graph abstraction. Diffusion pseudotime was subsequently calculated from the selected root using the same neighbour graph. For Palantir, the PCA embedding was used to compute diffusion maps and a multiscale diffusion representation, followed by pseudotime inference with the selected root supplied as the early cell.

Each non-degenerate pseudotime vector was independently min–max rescaled to $[0, 1]$ and aligned to the original cell order. Constant-valued outputs were assigned missing values, and method failures caught by the wrapper were represented by missing-value vectors. The resulting cell-by-method matrix provided a common input format for downstream BioTrajX evaluation. This standardisation retained a single ordering per method; lineage topology and fate probabilities were not included in the wrapper output.

#### LCMV Time-course CD8$^+$ T-cell Dataset (Ground-Truth Validation)(Figure S8)

Single-cell RNA sequencing data spanning a full time course of lymphocytic choriomeningitis virus (LCMV) infection were obtained from the NCBI Gene Expression Omnibus dataset GSE131847 [1], comprising CD8$^+$ T cells isolated at 12 time points post-infection (naive, d3, d4, d5, d6, d7, d10, d14, d21, d32, d60, d90). Because the day of isolation is recorded for every cell, this dataset provides direct ground truth against which a method's recovered pseudotime ordering can be validated.

Preprocessing followed the analysis code accompanying the review by Hudson and Wieland [2], deposited in Mendeley Data (2022, version 1; `https://doi.org/10.17632/3dvt79c7yt.1`). Raw count matrices from individual time points were merged into a single Seurat object (v. 5.0.2) and normalized using `SCTransform` (`vst.flavor = "v1"`). PCA was performed using 50 components, and UMAP was calculated from PCs 1–50.

For BioTrajX evaluation, the top 2,000 highly variable genes were selected from the SCT-normalized data, and both the HVG-restricted and all-gene log-normalized matrices were used as input to the trajectory-inference methods, as described above.

For the root-dependent methods, two root-selection configurations were evaluated within the naive population. The primary nearest-centroid root (NCR) was the observed naive cell closest to the naive-population centroid in PCA space. The alternative CytoTRACE root (CR) was the naive cell with the lowest inverted and rescaled CytoTRACE score, corresponding to the greatest estimated developmental potential. CytoTRACE pseudotime itself was computed once because it does not require an external root. Marker genes for the BioTrajX Directionality and Endpoint Validity scores were retrieved from MSigDB (*Mus musculus*) using `get_markers_msigdb()`: naive-pole markers were drawn from `GSE41867_NAIVE_VS_DAY8_LCMV_EFFECTOR_CD8_TCELL_UP` and effector-pole markers from `GSE41867_NAIVE_VS_DAY8_LCMV_EFFECTOR_CD8_TCELL_DN` (C7 immunological-signature collection), both filtered to the top 30 genes with detection in at least 10% of cells using `filter_markers()`.

**Trajectory evaluation and recovery of sampling time** Pseudotimes generated by the eight-method wrapper were evaluated using the SCT assay of the processed Seurat object. Methods with more than 10 non-missing pseudotime values were retained for evaluation with `compute_multi_doe_linear`, using the filtered early and terminal marker sets described above. Endpoint Validity was calculated with the Gaussian mixture model option (`E_method = "gmm"`); other scoring parameters were left at their defaults. The component scores and composite DOE score were summarised in a heatmap, and methods were ordered by decreasing DOE score for the pseudotime UMAP and marker-module trend displays.

Early and terminal module scores were calculated separately using Seurat's `AddModuleScore`. Cells were matched to the pseudotime matrix by cell identifier, and cells with missing pseudotime were excluded separately for each method. Module scores were plotted against pseudotime with locally estimated scatterplot smoothing (LOESS; span = 0.4) and the default 95% confidence bands.

Recorded sampling times provided an external temporal reference. The naive group was assigned day 0, and the remaining groups were assigned their corresponding sampling days: 3, 4, 5, 6, 7, 10, 14, 21, 32, 60, and 90. For each method with at least 10 matched cells with available pseudotime, Spearman and Pearson correlations between pseudotime and sampling day were calculated using `cor.test`. Signed Spearman correlation was used as the primary measure of temporal-order recovery. Its association with DOE score was then assessed across methods using both Spearman and Pearson correlations, provided that at least three methods were available. Pseudotime distributions within each sampling group were visualised using violin plots with overlaid boxplots, with methods ordered by decreasing Spearman correlation with sampling day.

#### CD8$^{+}$ T-cell Exhaustion Dataset (Linear Trajectory)(Figure S9)

Single-cell RNA sequencing data from head and neck squamous cell carcinoma (HNSCC) patients were obtained from the NCBI Gene Expression Omnibus dataset GSE164690 [3], comprising matched peripheral blood lymphocyte (PBL) and CD45$^{+}$ tumor-infiltrating lymphocyte (TIL) samples from 17 patients (HN01–HN18, excluding HN10, for which no matched PBL sample was available). Raw count matrices were loaded into Seurat (v. 5.0.2) and subjected to quality-control filtering, retaining cells with 200–3,000 detected genes (`nFeature_RNA`) and less than 10% mitochondrial reads (`percent.mt`).

Filtered cells were log-normalized using `NormalizeData`, and the top 2,000 highly variable genes were identified using variance-stabilizing transformation (VST) prior to data scaling. CD8$^{+}$ T cells were isolated by requiring co-expression of *CD3D*, *CD3G*, *CD3E*, *CD8A*, and *CD8B*. Functional states of CD8$^{+}$ T cells were annotated using ProjecTILs with the `CD8T_human_ref_v1` human reference atlas. Each cell was assigned a functional cluster label spanning four cell states: `CD8.NaiveLike`, `CD8.TPEX` (progenitor exhausted), and `CD8.TEX` (terminally exhausted). The expected biological trajectory is approximately linear, progressing from naive-like progenitors through memory and progenitor-exhausted states toward terminal exhaustion.

The top 2,000 highly variable genes were selected, and a log-normalized genes-by-cells expression matrix was used as input to the trajectory-inference methods. Early markers were drawn from the `CD8+ Tn` signature in the SlimR implementation of the curated pan-cancer tumor-infiltrating T-cell atlas [4], whereas terminal markers were retrieved from MSigDB using `get_markers_msigdb()` and drawn from the `GSE9650_EFFECTOR_VS_EXHAUSTED_CD8_TCELL_DN` gene set. Both marker sets were filtered to genes detected in at least 10% of cells using `filter_markers()`.

Table 3: Transcription factors used to validate pseudotime trends in the CD8+T exhaustion in cancer (Figure S9) .

| TF | Expected stage | Biological role | Reference |
|---|---|---|---|
| *TCF7* (*TCF1*) | Naive / stem-like (early) | Key regulator of stem-like and progenitor CD8$^{+}$ T-cell states; supports persistence and proliferative capacity while opposing terminal differentiation. | Siddiqui et al. [5] Rong et al. [6] Rangel Rivera et al. [7] |
| *LEF1* | Naive / stem-like (early) | Wnt-pathway TF associated with less differentiated, memory-like, and stem-like CD8$^{+}$ T-cell states and maintenance of T-cell persistence. | Zhou and Xue [8] Rangel Rivera et al. [7] Miakicheva et al. [9] |
| *FOXP1* | Naive / quiescent (early) | Transcriptional regulator associated with T-cell quiescence and restrained effector differentiation in less differentiated CD8$^{+}$ T-cell states. | Stephen et al. [10] Zhu et al. [11] Wizzard et al. [12] |
| *TOX* | Exhausted (terminal) | Central regulator of CD8$^{+}$ T-cell exhaustion that establishes and maintains exhaustion-associated transcriptional and epigenetic programs. | Khan et al. [13] Kim et al. [14] Huang et al. [15] |
| *BATF* | Exhaustion-associated (terminal) | AP-1 family TF associated with chronic T-cell stimulation and regulation of exhaustion-associated transcriptional programs in tumour-reactive CD8$^{+}$ T cells. | Zhang et al. [16] |
| *EOMES* | Differentiated / exhausted (terminal) | T-box TF associated with differentiated and dysfunctional CD8$^{+}$ tumour-infiltrating T-cell states and exhaustion-related transcriptional programs. | Li et al. [17] Weulersse et al. [18] Llaó-Cid et al. [19] |

**Trajectory evaluation and transcription-factor trends** For BioTrajX evaulation, pseudotimes generated by the unified wrapper were evaluated using `compute_multi_doe_linear` with the filtered naive and terminal marker sets described above. Methods with more than 10 non-missing pseudotime values were retained. Endpoint Validity was calculated using `E_method = "gmm"`, with other scoring parameters left at their defaults. Component scores and composite DOE scores were displayed in a heatmap, and methods were ranked by decreasing DOE score.

Pseudotimes were overlaid on the existing Seurat UMAP embedding alongside the functional-state annotations. Early and terminal module scores were calculated separately using `AddModuleScore` and plotted against each method's pseudotime using LOESS smoothing with span = 0.4 and default 95% confidence bands. Cells with missing pseudotime were excluded separately for each method.

To inspect transcriptional patterns beyond the module-level summaries, normalised expression values were extracted for the early-associated transcription factors *TCF7*, *LEF1*, and *FOXP1*, and the terminal-associated transcription factors *TOX*, *BATF*, and *EOMES* (Table 3). For each available gene and TI method, expression was modelled as a smooth function of pseudotime using a generalised additive model with the formula `y ~ s(x, bs = "cs")`, specifying a shrinkage cubic regression spline. Fitted curves and default 95% confidence bands were displayed alongside cell-level expression values. These fits provided descriptive assessments of gene-expression trends.

Pseudotime distributions were also compared across the annotated functional states using violin plots with overlaid boxplots. States were displayed in the order CD8.NaiveLike, CD8.EM, CD8.TPEX, and CD8.TEX, and method panels followed the DOE ranking.

#### Hematopoietic Stem Cell Differentiation Dataset (Branched Trajectory)(Figure S10)

The ageing haematopoietic stem cell dataset, originally generated by Kowalczyk et al. [20], was obtained as a preprocessed and batch-corrected expression matrix from the CytoTRACE website [21]. The dataset comprises long-term HSCs, short-term HSCs, and multipotent progenitors isolated from young and old mice.

The trajectory is bifurcated, with a shared stem/progenitor compartment (Stem_Progenitors) diverging into an erythroid lineage (Erythroid_progenitors_Erythroblasts, Erythrocytes) and a B-cell lineage (Immature_B, Mature_B).

BioTrajX evaluated the two lineages independently as separate branches: Stem→Erythroid and Stem→B cell. Early markers shared across both branches were drawn from the `HAY_BONE_MARROW_CD34_POS_HSC` gene set (C8 cell type signatures; Human Cell Atlas bone marrow $CD34^+$ HSC signature, ortholog-mapped to mouse). Erythroid terminal markers were drawn from the `WP_ERYTHROPOIESIS` gene set (C2 WikiPathways curated erythropoiesis pathway), and B-cell terminal markers were drawn from the `HADDAD_B_LYMPHOCYTE_PROGENITOR` gene set (C2 curated gene sets). All marker sets were filtered to the top 30 genes with at least 10% detection in the respective Seurat object.

**Branch-specific trajectory evaluation** The pseudotime vectors produced by the unified wrapper were evaluated separately within two branches defined by the Seurat object's `Phenotype` annotations. The Stem-to-Erythroid branch included `Stem_Progenitors`, `Erythroid_progenitors_Erythroblasts`, and `Erythrocytes`; the Stem-to-B-cell branch included `Stem_Progenitors`, `Immature_B`, and `Mature_B`. Thus, the stem/progenitor compartment was included in both branches. Branch membership was specified from these annotations rather than inferred from each method's trajectory topology.

Methods with more than 10 non-missing pseudotime values across the dataset were supplied to `compute_multi_doe_branched`. Branch filters specified a minimum of 50 cells, and each branch was evaluated using the shared early-marker set and its corresponding terminal-marker set. Marker filtering was performed on the full Seurat object before branch-specific evaluation. Endpoint Validity used `E_method = "gmm"`, with other scoring parameters left at their defaults. Branch-specific component scores, branch-specific DOE scores, and the aggregate comparison returned by BioTrajX were recorded. The existing full-dataset pseudotime vectors were supplied for branch evaluation; TI methods were not rerun separately within each branch.

Whole-dataset UMAP displays were ordered by aggregate DOE score, whereas branch-specific displays were ordered independently by the corresponding branch's DOE score. For branch-specific UMAP

Table 4: Transcription factors used to validate pseudotime trends in the haematopoietic stem-cell differentiation trajectories (Figure S10).

| TF | Expected stage | Biological role | Reference |
|---|---|---|---|
| *Hlf* | Stem/progenitor (early) | PAR-bZIP TF enriched in haematopoietic stem cells that promotes quiescence, self-renewal, and maintenance of stem-cell potential. | Komorowska et al. [22]<br>Lee et al. [23]<br>Wu et al. [24] |
| *Meis1* | Stem/progenitor (early) | TALE-homeodomain TF associated with HSC quiescence and long-term maintenance, including regulation of metabolic and oxidative-stress programs. | Kocabas et al. [25]<br>Unnisa et al. [26]<br>Coulombe et al. [27] |
| *Klf1* (*EKLF*) | Erythroid lineage (terminal) | Major erythroid TF coordinating lineage-specific gene expression, globin production, and terminal erythroid differentiation. | Tallack et al. [28]<br>Mukherjee et al. [29]<br>Mukherjee and Bieker [30] |
| *Nfe2* | Erythroid lineage (terminal) | Erythroid TF associated with late erythroid maturation and regulation of globin expression, heme biosynthesis, and erythroid gene programs. | Shivdasani and Orkin [31]<br>Rheinemann et al. [32]<br>Gothwal et al. [33] |
| *Bach2* | B-cell lineage (late) | Transcriptional regulator of B-cell differentiation that maintains B-cell identity and restrains premature plasma-cell differentiation. | Muto et al. [34]<br>Itoh-Nakadai et al. [35]<br>Shinnakasu et al. [36] |
| *Ikzf3* (*Aiolos*) | B-cell lineage (late) | Ikaros-family TF associated with B-cell maturation and regulation of B-cell activation, identity, and receptor-signalling programs. | Wang et al. [37]<br>Lazarian et al. [38]<br>Yamashita and Morio [39] |

colouring, pseudotime was additionally min–max rescaled within each displayed branch. Module-trend, transcription-factor, and violin plots retained the pseudotime values from the original wrapper output.

**Branch-specific expression trends** Early, erythroid, and B-cell module scores were calculated on the full Seurat object using `AddModuleScore`. Within each branch, the early module and the corresponding terminal module were plotted against pseudotime using LOESS smoothing with span = 0.5 and default 95% confidence bands.

Transcription-factor trends were examined separately for each branch using normalised expression values. Both branches included the early-associated factors *Hlf* and *Meis1*; terminal-associated factors were *Klf1* and *Nfe2* for the erythroid branch and *Bach2* and *Ikzf3* for the B-cell branch (Table 4). For each available gene and method, generalised additive models were fitted using `y ~ s(x, bs = "cs")`, with fitted curves and default 95% confidence bands displayed. Cells with missing pseudotime were excluded separately for each method. Violin plots with overlaid boxplots summarised pseudotime distributions across the three annotated cell populations within each branch.

#### Root cell selection for Monocle3 in Supplementary (Figure S11)

To assess BioTrajX as an aid for comparing candidate trajectory roots, seven root-selection configurations were defined on the $CD8^+$ T-cell exhaustion dataset before running Monocle3. Three candidates were selected from the CD8.NaiveLike population: the cell with the lowest inverted CytoTRACE score, cell with the lowest module score calculated from the naive marker sets; the cell nearest the CD8.NaiveLike centroid in PCA space; and the cell with the highest naive-marker module score, calculated using `AddModuleScore`. The naive-marker set was the same filtered `KAECH_NAIVE_VS_DAY8_EFF_CD8_TCELL_UP` signature used for Figure S9.

The remaining three configurations used centroid-representative cells from the CD8.EM, CD8.TPEX, and CD8.TEX clusters as alternative roots. For each cluster, the candidate root was selected using the same centroid-based procedure described above. Together with the CD8.NaiveLike root, this yielded five root configurations.

Monocle3 was run separately for each root-selection configuration using the same expression matrix restricted to the top 2,000 highly variable genes. Each run repeated preprocessing, UMAP construction, clustering, principal-graph learning with `use_partition = FALSE`, and cell ordering from the selected root. Infinite pseudotimes were converted to missing values, and the resulting pseudotime vector was min–max rescaled to $[0, 1]$.

Each resulting pseudotime vector was evaluated using `compute_multi_doe_linear`, with the same filtered early and terminal marker sets as Figure S9 and `E_method = "gmm"`. Composite DOE scores were compared across root-selection configurations, while radar plots and heatmaps displayed the component scores. Pseudotimes and candidate-root positions were overlaid on the existing Seurat UMAP embedding. Naive and terminal module scores were plotted against pseudotime using LOESS smoothing with span = 0.4 and default 95% confidence bands.

### Supplementary Results S4. Synthetic Dataset Simulations

**Figure S1 DOE scores across linear and branched synthetic datasets** Heatmaps summarise $D_{\text{early}}$, $D_{\text{term}}$, $O$, $E_{\text{early}}$, $E_{\text{term}}$, and the composite DOE score. **(A)** Linear datasets representing correctly oriented, reversed, stepwise, zero-inflated negative binomial, flat, high-dropout, and low-signal trajectories. **(B)** Corresponding branched datasets evaluated along the $A \to B$ lineage. The sign shown with $O$ indicates whether the orientation selected by Order Consistency agrees with the expected early-to-terminal direction.

**Figure S2 UMAP visualisation of linear and branched synthetic datasets** UMAP embeddings show the expression-space structure of the synthetic datasets, with cells coloured by the pseudotime assigned in each dataset. **(A)** Gaussian linear; **(B)** Gaussian reversed; **(C)** stepwise repression; **(D)** stepwise activation; **(E)** ZINB linear; **(F)** ZINB reversed; **(G)** flat/no signal; **(H)** high dropout; **(I)** low signal; **(J)** Gaussian branched; **(K)** Gaussian branched reversed; **(L)** branched stepwise repression; **(M)** branched stepwise activation; **(N)** branched ZINB; **(O)** branched low signal; **(P)** ZINB linear with shuffled pseudotime; and **(Q)** branched ZINB with shuffled pseudotime. Branch identities are indicated for the branched datasets.

**Figure S3 Marker-gene dynamics across synthetic datasets** Expression profiles of representative early, terminal, and background genes are plotted against pseudotime. Points represent individual cells and smoothed curves summarise gene-expression trends. Panels **A**–**Q** correspond to the synthetic scenarios shown in Figure S2; branched panels display the $A \to B$ lineage. Correctly oriented datasets show decreasing early-marker expression and increasing terminal-marker expression, whereas reversed, flat, dropout, and low-signal datasets illustrate distinct departures from the expected progression.

**Figure S4 Directionality diagnostics across synthetic datasets** Per-cell early and terminal marker scores are plotted against pseudotime for each synthetic scenario. Spearman correlations ($\rho$) quantify monotonic association, and the corresponding ReLU-transformed Directionality scores are displayed above each plot. Panels **A**–**Q** follow the dataset order in Figure S2. Correctly oriented trajectories produce positive early- and terminal-direction scores, whereas reversed trajectories are assigned scores of zero.

**Figure S5 Order Consistency diagnostics across synthetic datasets** Representative marker-gene expression profiles are shown against pseudotime after orientation according to the expected direction of marker change. Coloured step functions indicate direction-constrained isotonic regression fits. The null-calibrated Order Consistency score and selected orientation are reported in each panel. Panels **A**–**Q** follow the dataset order in Figure S2. Reversed trajectories retain high $O$ scores but receive a negative orientation sign, whereas flat and weak-signal datasets show reduced monotonic consistency.

**Figure S6 Endpoint Validity diagnostics across synthetic datasets** Precision-at-$k$ curves show the enrichment of expected early cells among cells with the lowest pseudotime and expected terminal cells among cells with the highest pseudotime. Blue and red curves represent early and terminal endpoint recovery, respectively. Filled points indicate the $k$ values used to calculate $E_{\text{early}}$ and $E_{\text{term}}$, and the dotted horizontal line indicates the corresponding endpoint prevalence. Panels **A**–**Q** follow the dataset order in Figure S2.

**Figure S7 Recovery of known endpoints in stepwise synthetic datasets** Marker-score-derived endpoint labels obtained using Gaussian mixture modelling are compared with the known endpoint states of the synthetic datasets. Cells are classified as true positives, false negatives, false positives, or true negatives separately for the early and terminal endpoints. **(A)** Stepwise repression; **(B)** reversed stepwise activation; **(C)** branched stepwise repression along the $A \to B$ lineage; and **(D)** reversed branched stepwise activation along the $A \to B$ lineage. Correctly oriented datasets recover the known endpoints with high Endpoint Validity, whereas reversed datasets place the endpoint populations at the incorrect pseudotime extremes.

**Figure S1. Summary heatmap of DOE scores across synthetic fixtures**

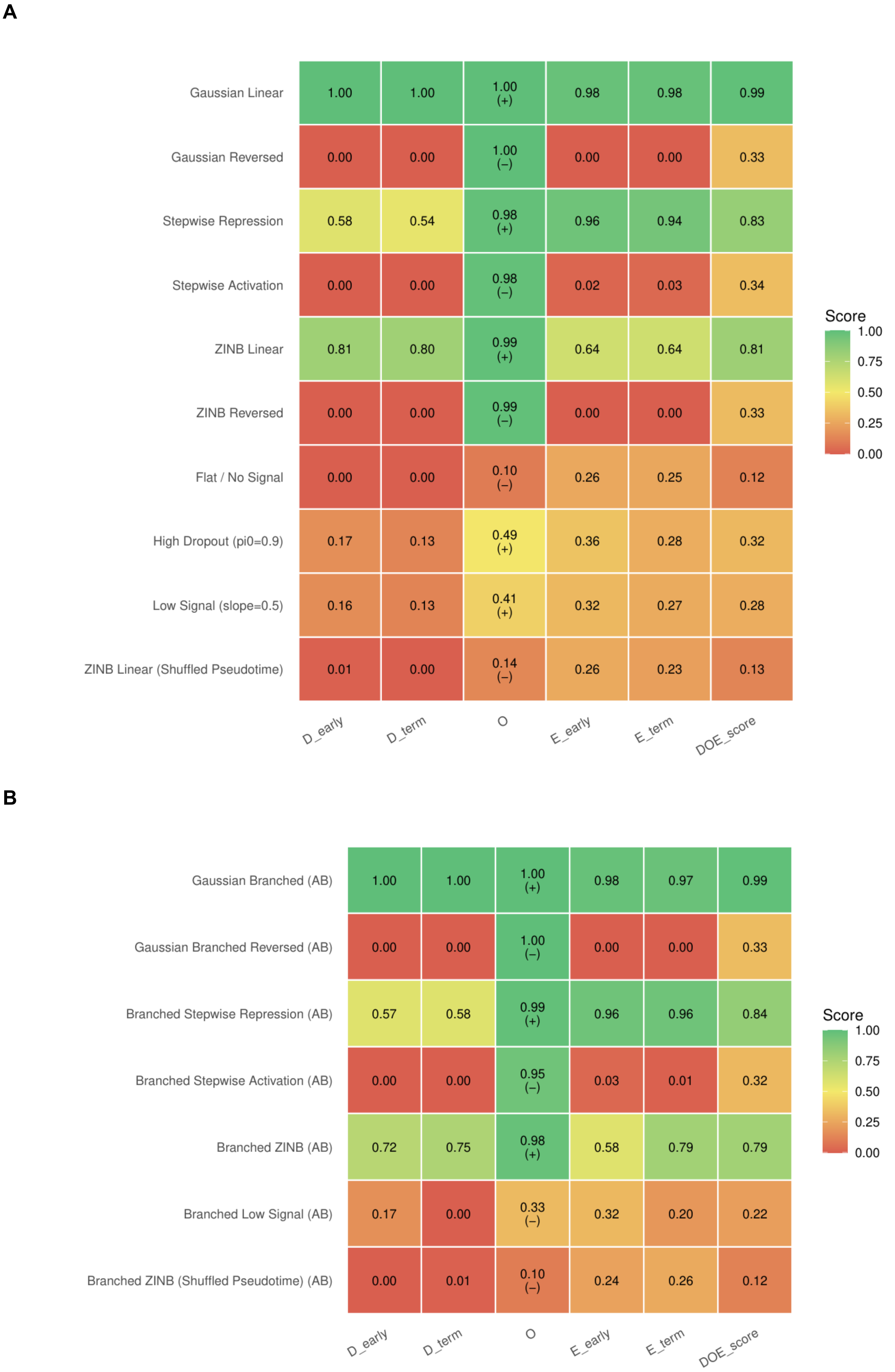

### Figure S2. UMAP embeddings of synthetic trajectory fixtures

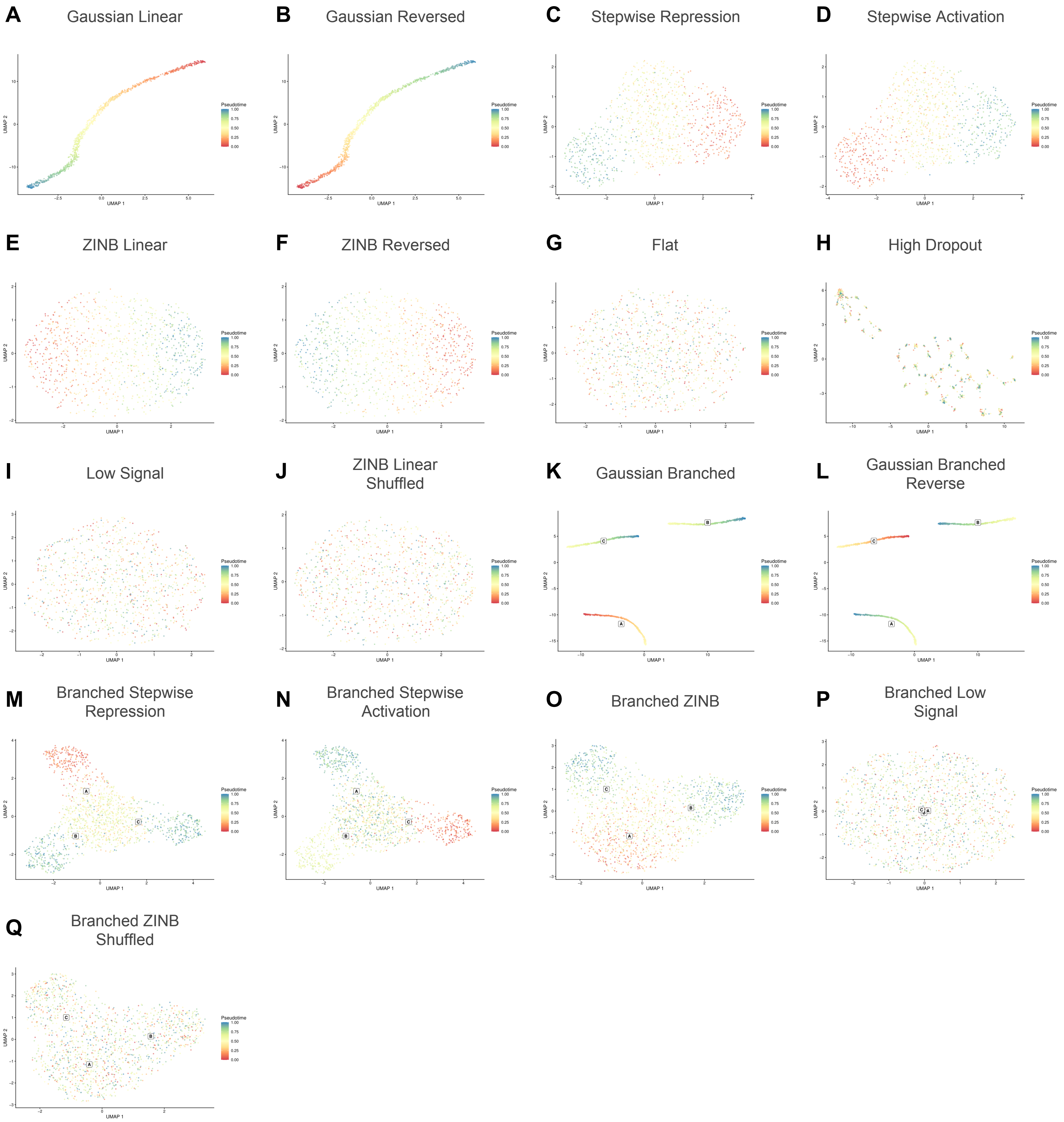

#### Figure S3. Gene expression patterns across synthetic fixtures

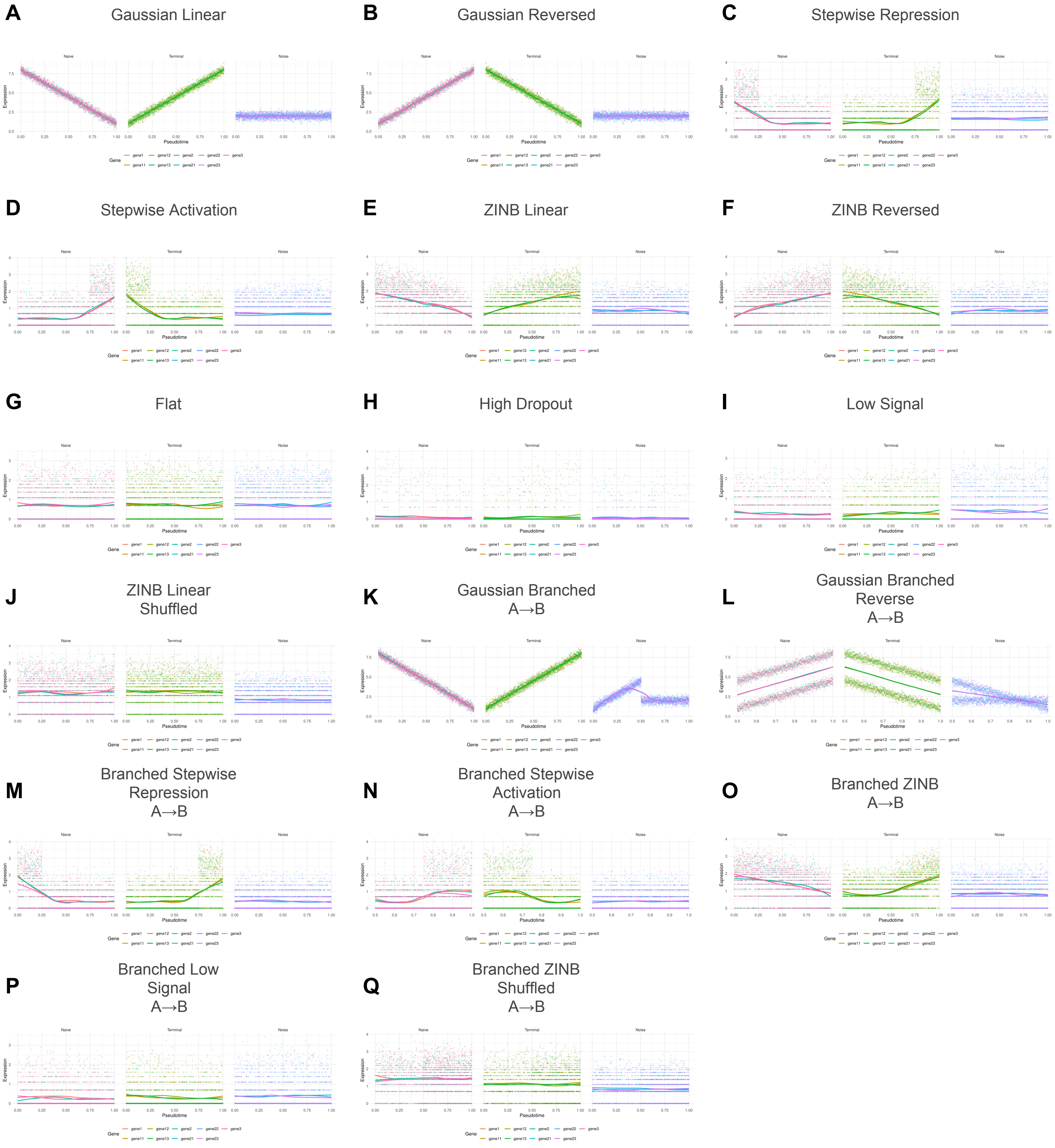

### Figure S4. Directionality (D) metric evaluation across synthetic fixtures

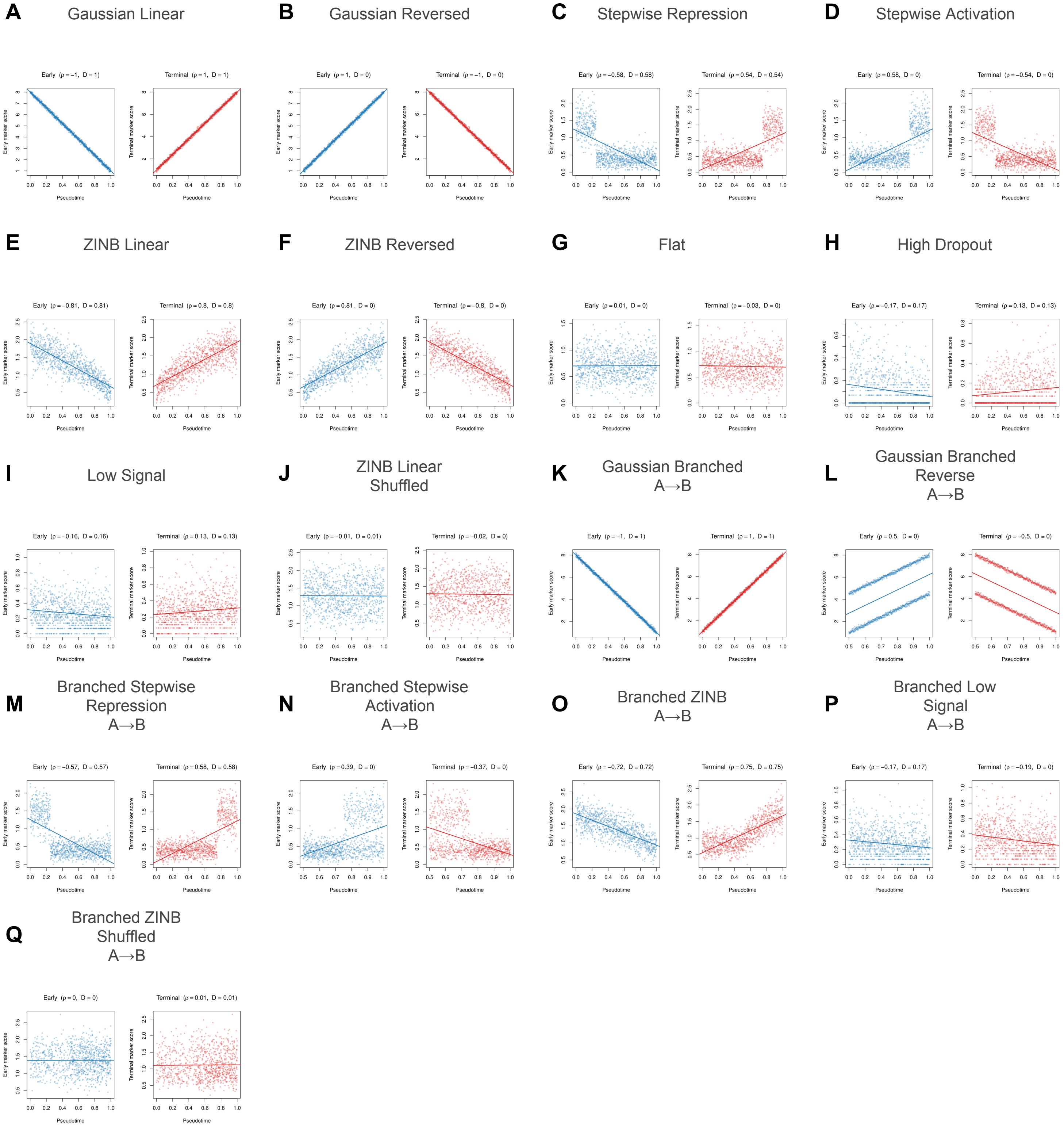

#### Figure S5. Order consistency (O) evaluation across synthetic fixtures

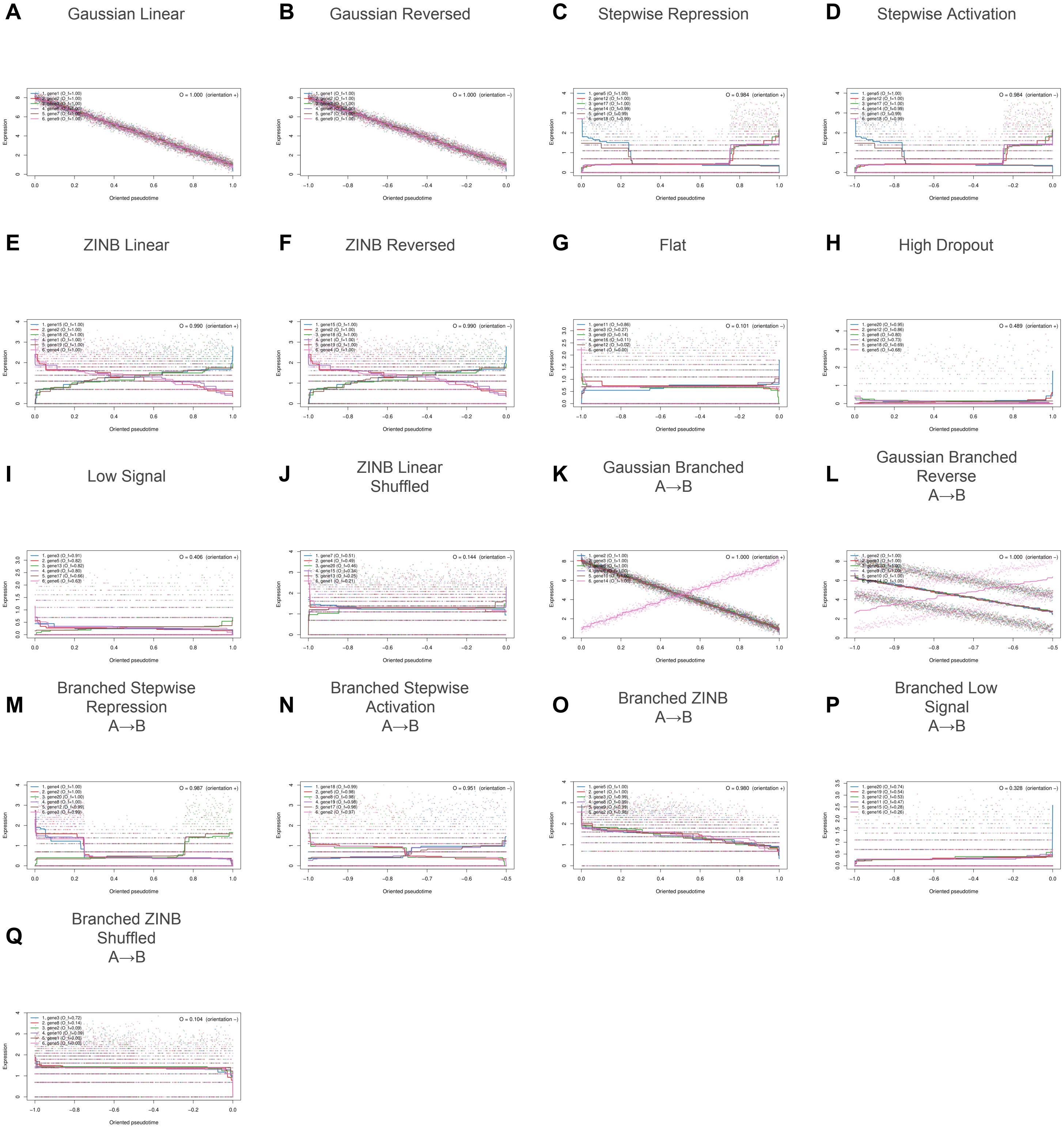

#### Figure S6. Endpoint validity (E) evaluation across synthetic fixtures

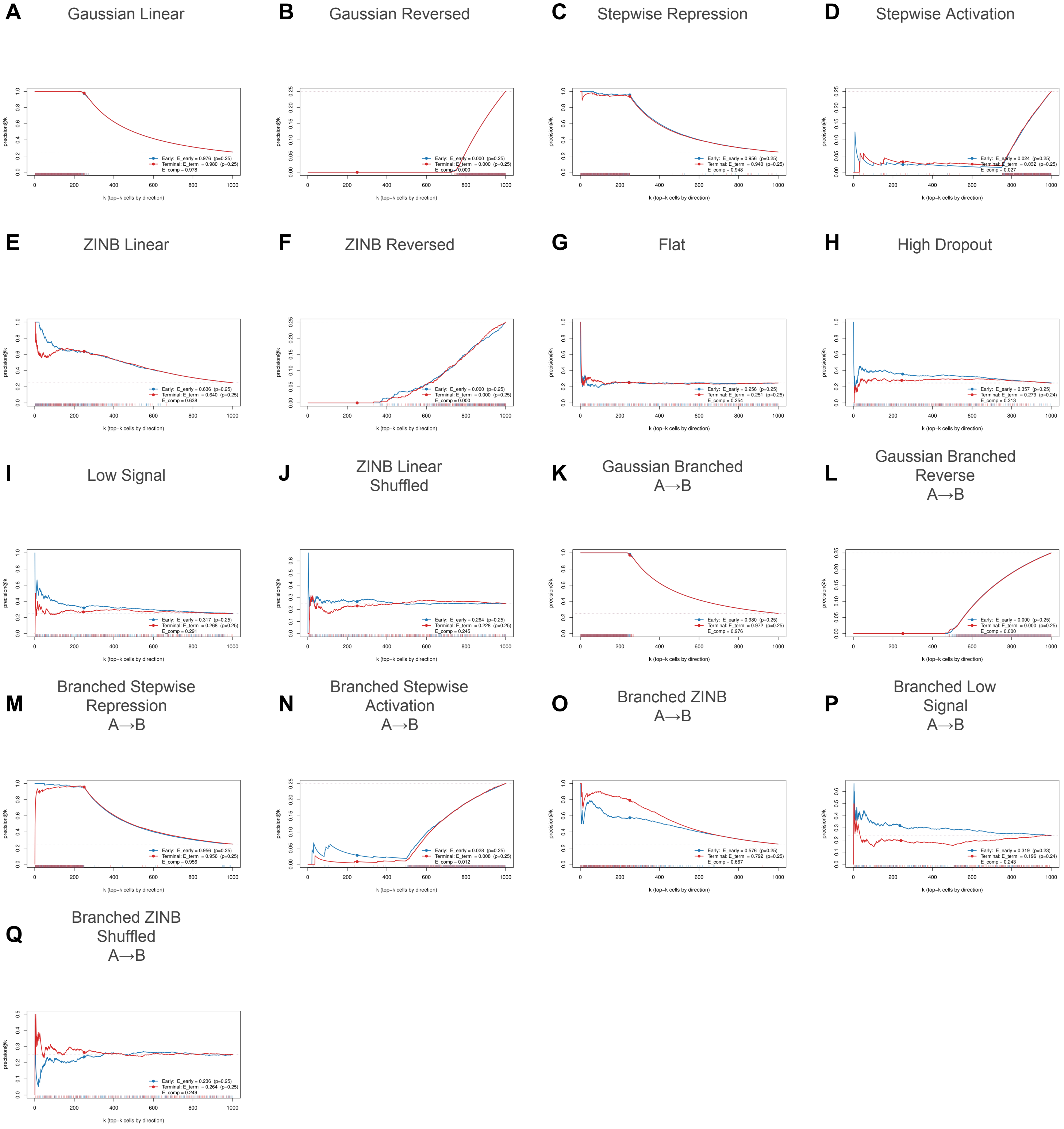

### Figure S7. Ground-truth endpoint recovery across stepwise and branched stepwise fixtures

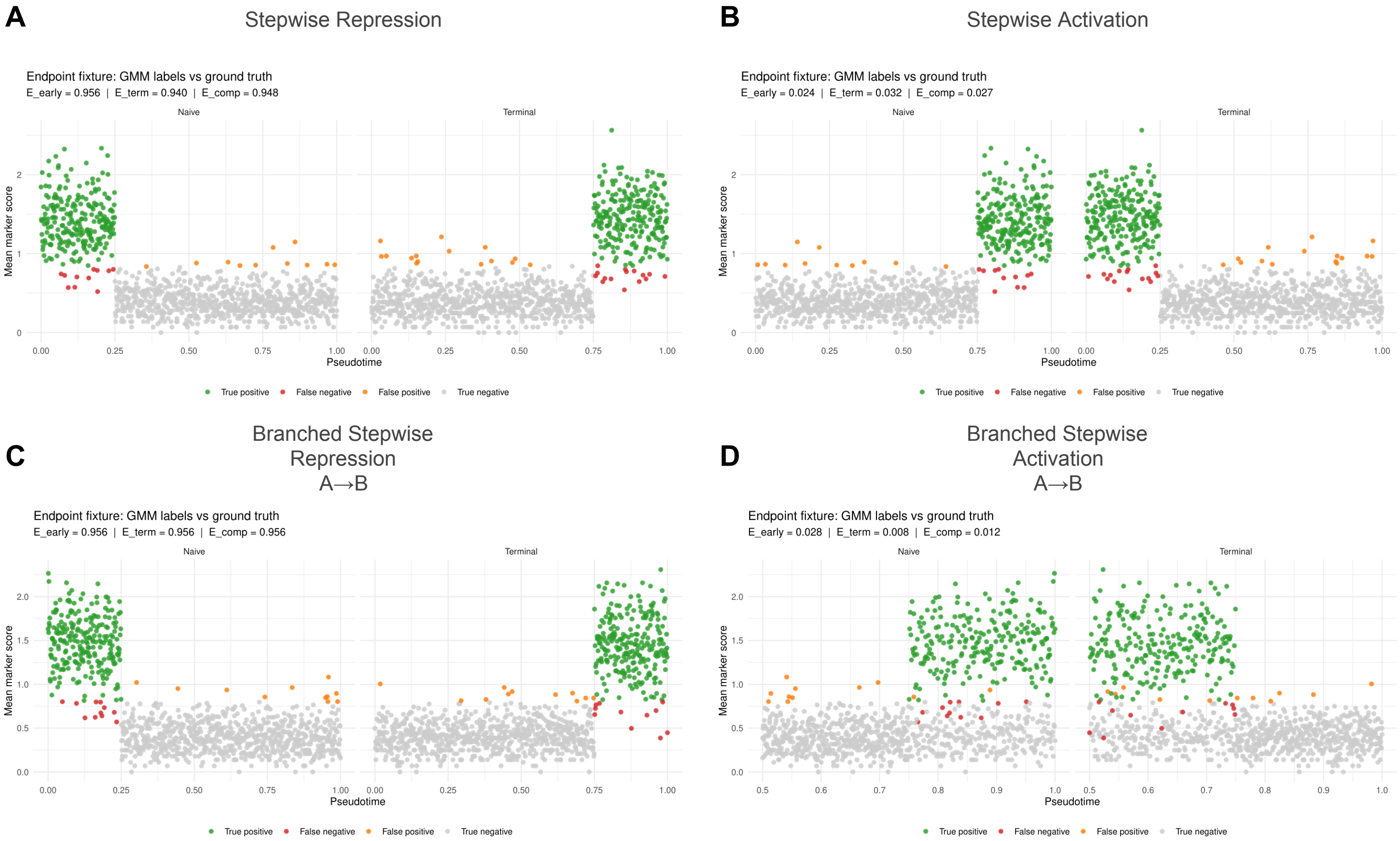

### Supplementary Results S5. Real Dataset Applications

**Figure S8 BioTrajX DOE score ranking correlates with ground-truth infection timing recovery (GSE131847)** **(A)** UMAP embedding coloured by experimentally observed day post-infection and by pseudotime inferred using CytoTRACE and seven root-dependent trajectory-inference methods evaluated using two root-selection strategies. CR denotes the CytoTRACE-selected root and NCR denotes the naive-centroid root. The composite DOE score is shown above each trajectory. **(B)** Smoothed early and terminal marker-module scores along the pseudotime inferred by each method. **(C)** Heatmap of DOE component and composite scores across methods. The sign shown with $O$ indicates the inferred trajectory orientation, and the boxed final column contains the composite DOE score. **(D)** Distribution of inferred pseudotime across experimental days for each method, showing differences in recovery and orientation of the known temporal progression.

**Figure S9 BioTrajX DOE evaluation of CD8$^+$ T-cell exhaustion as a linear trajectory** **(A)** UMAP embedding coloured by annotated CD8$^+$ T-cell state and by pseudotime inferred using eight trajectory-inference methods. The composite DOE score is shown above each method-specific embedding. **(B)** Smoothed early and terminal marker-module scores along the pseudotime inferred by each method. **(C)** Heatmap of DOE component and composite scores across methods. The sign shown with $O$ indicates the inferred trajectory orientation, and the boxed final column contains the composite DOE score. **(D)** Smoothed expression trajectories of representative exhaustion-associated transcription factors along inferred pseudotime, grouped into early-state TFs (*TCF7*, *LEF1*, and *FOXP1*) and terminal/exhaustion-associated TFs (*TOX*, *BATF*, and *EOMES*). **(E)** Distribution of inferred pseudotime across annotated CD8$^+$ T-cell states for each trajectory configuration.

**Figure S10 Branch-specific BioTrajX DOE evaluation of haematopoietic stem-cell differentiation** **(A)** UMAP embedding coloured by annotated cell type and by pseudotime inferred using eight trajectory-inference methods. **(B)** Heatmaps of DOE component and composite scores for the Stem-to-B and Stem-to-Erythroid branches, together with an overall DOE score summarising performance across both branches. **(C)** Branch-specific UMAP embeddings coloured by annotated developmental state and inferred pseudotime for the erythroid and B-cell branches, with methods ordered by branch-specific DOE score. **(D)** Smoothed early and lineage-specific terminal marker-module scores along inferred pseudotime for each branch. **(E–F)** Smoothed expression trajectories of representative lineage-associated transcription factors along inferred pseudotime for the erythroid (**E**) and B-cell (**F**) branches. Early-state TFs are *Hlf* and *Meis1*; lineage-associated terminal TFs are *Klf1* and *Nfe2* for the erythroid branch and *Bach2* and *Ikzf3* for the B-cell branch. **(G–H)** Distribution of inferred pseudotime across annotated developmental stages for the erythroid (**G**) and B-cell (**H**) branches, illustrating branch-specific differences in recovery of the expected developmental progression.

**Figure S11 BioTrajX-guided comparison of Monocle3 root-cell choices in CD8$^+$ T-cell exhaustion** **(A)** UMAP embedding coloured by annotated CD8$^+$ T-cell state and by Monocle3 pseudotime obtained using six candidate root cells. Black triangles indicate the selected roots. Candidates comprise three cells from the CD8.NaiveLike population—the cell with the lowest inverted CytoTRACE score, the cluster-centroid cell, and the cell with the highest naive-marker-module score—and the centroid cells from the CD8.EM, CD8.TPEX, and CD8.TEX populations. The composite DOE score is shown above each trajectory. **(B)** Smoothed early and terminal marker-module scores along pseudotime for each root initialization. **(C)** Composite DOE scores ranked by candidate root, with roots from the CD8.NaiveLike population distinguished from those in later cell states. **(D)** Radar-plot comparison of DOE components across root initializations.

### Figure S8. BioTrajX DOE score ranking correlates with ground-truth infection timing recovery

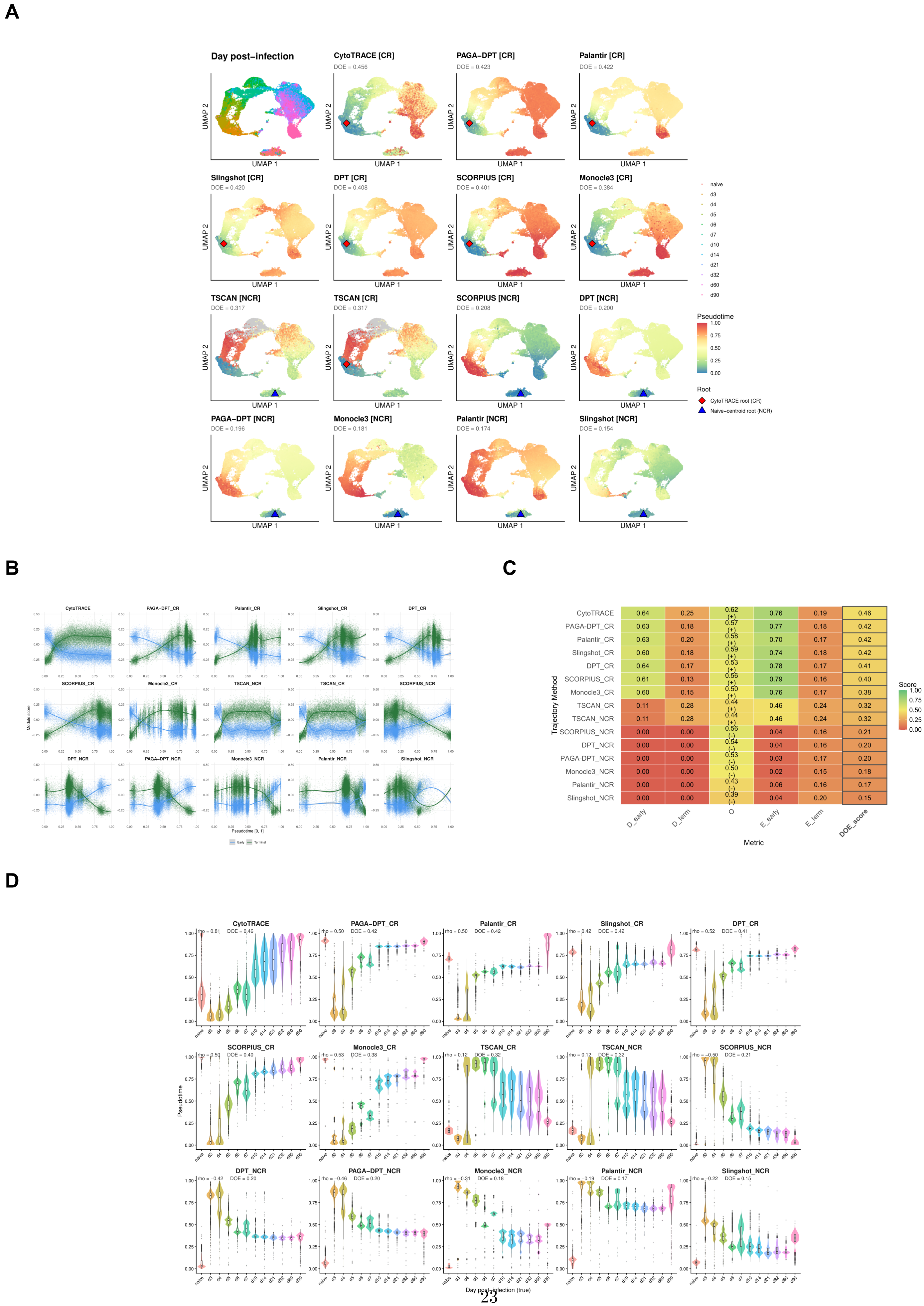

### Figure S9. BioTrajX DOE evaluation on CD8+ T cell exhaustion (linear trajectory)

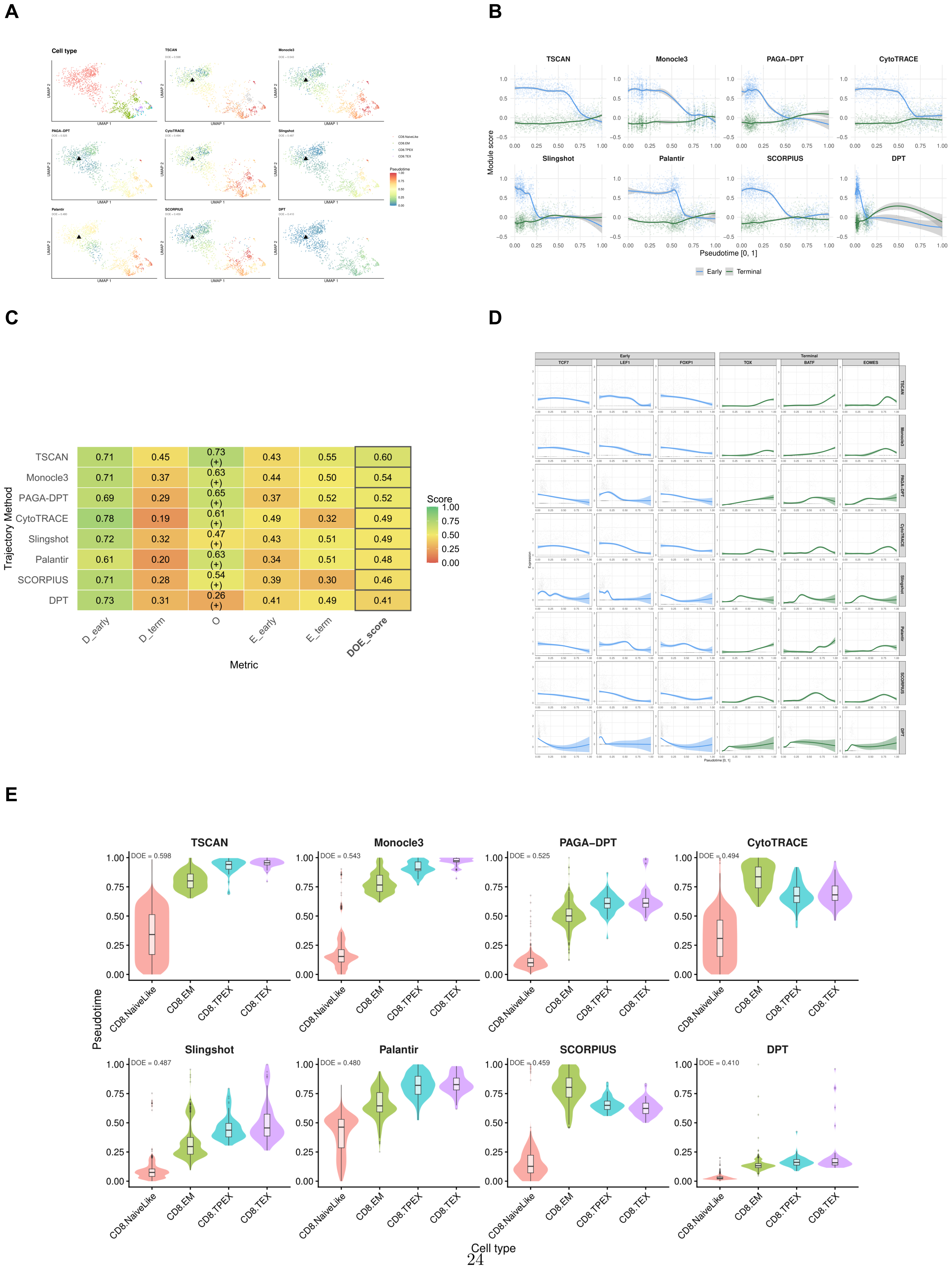

### Figure S10. BioTrajX DOE evaluation on stem cell differentiation (branched trajectory)

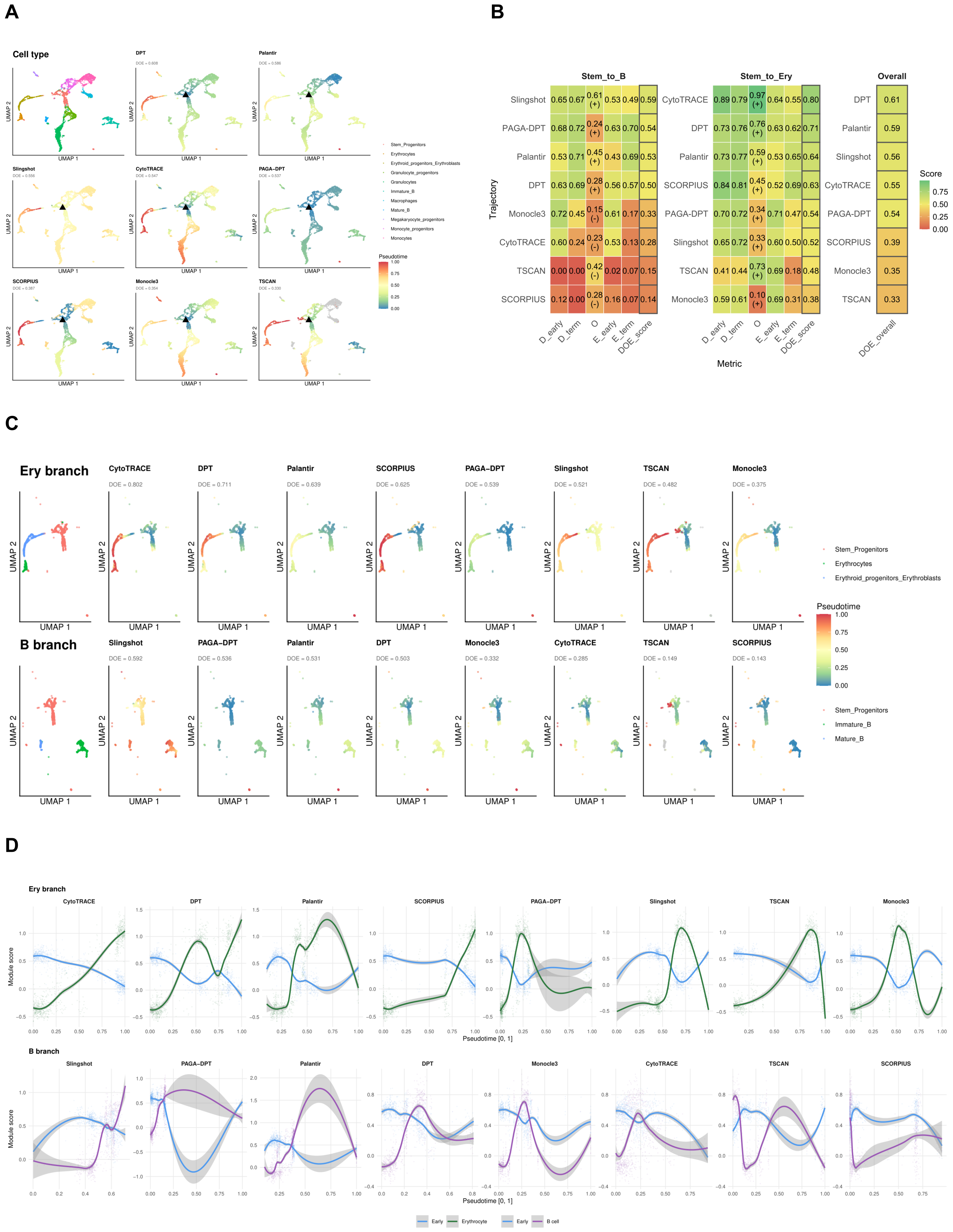

### Figure S10 (continued). BioTrajX DOE evaluation on stem cell differentiation (branched trajectory)

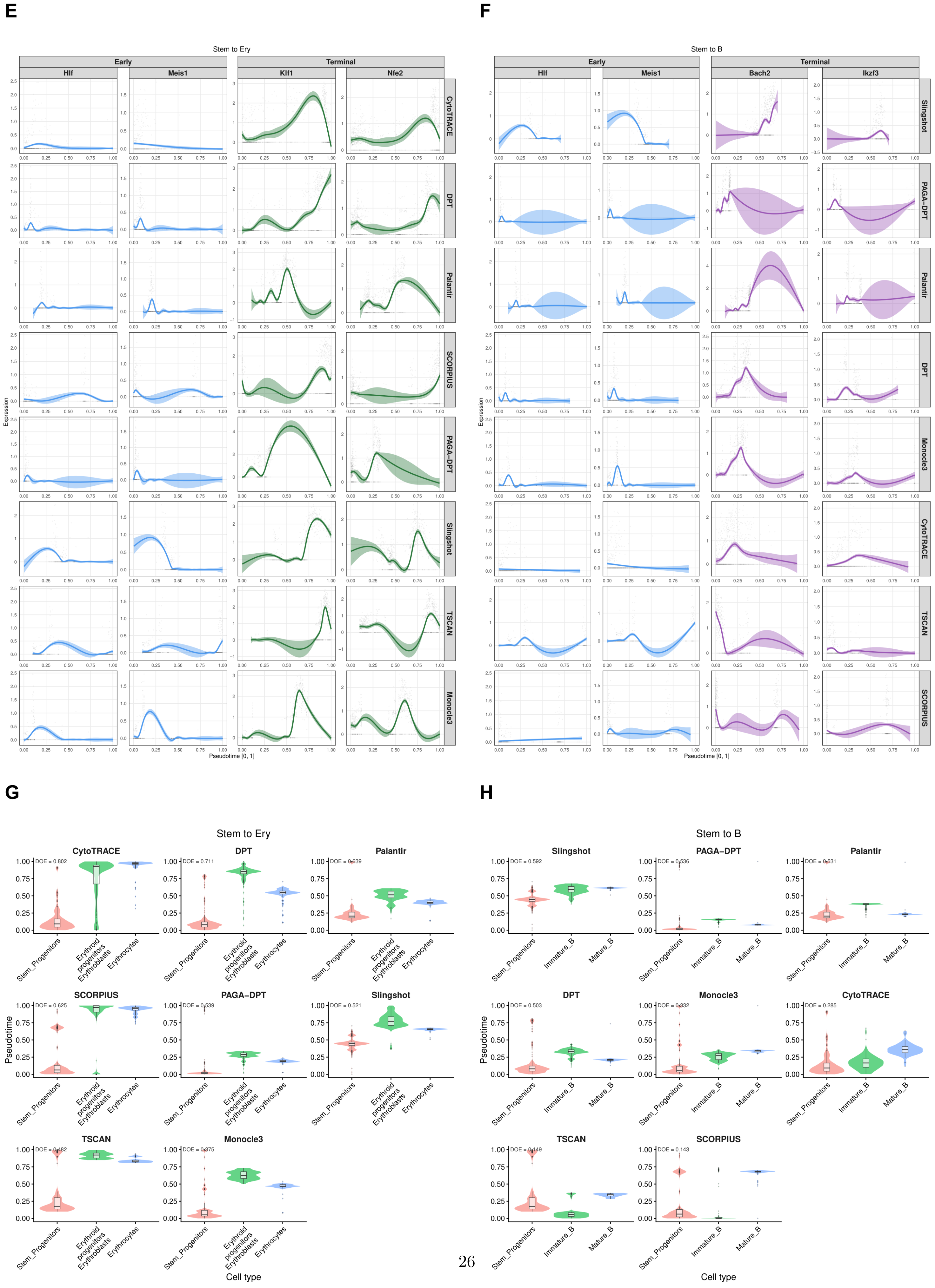

**Figure S11. BioTrajX DOE-guided root selection with Monocle3 on CD8+ T cells**

A

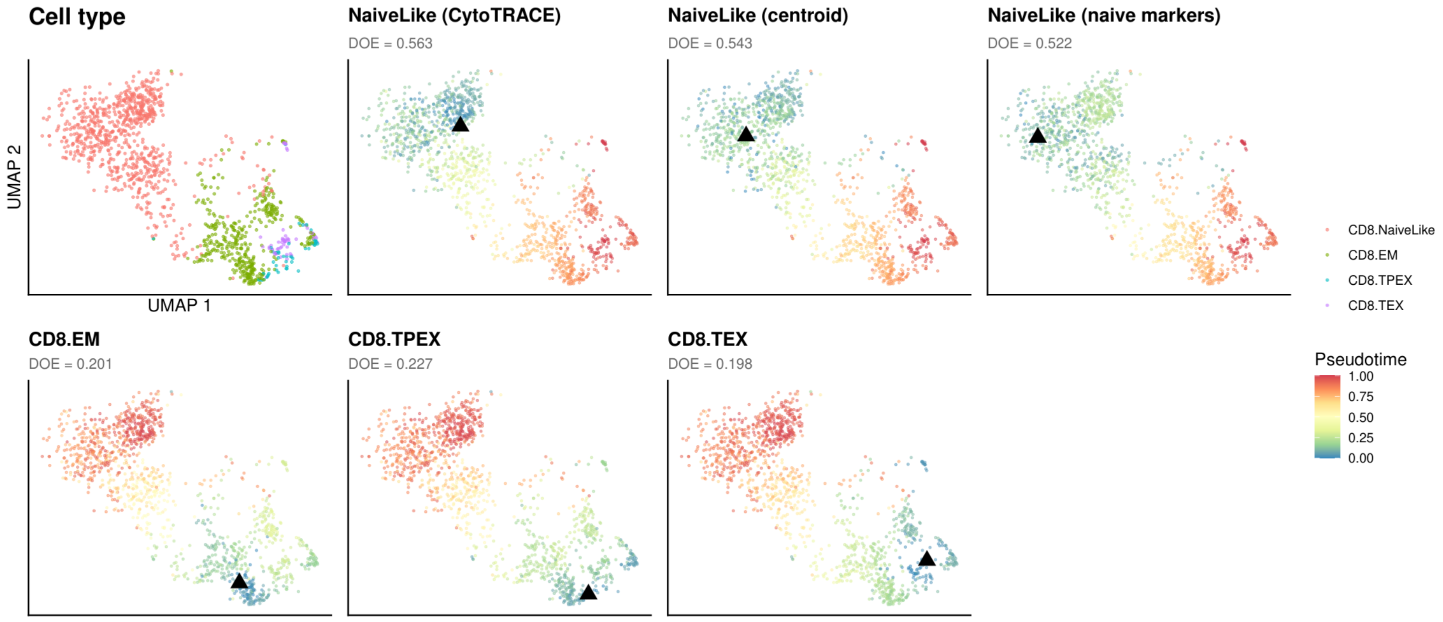

B

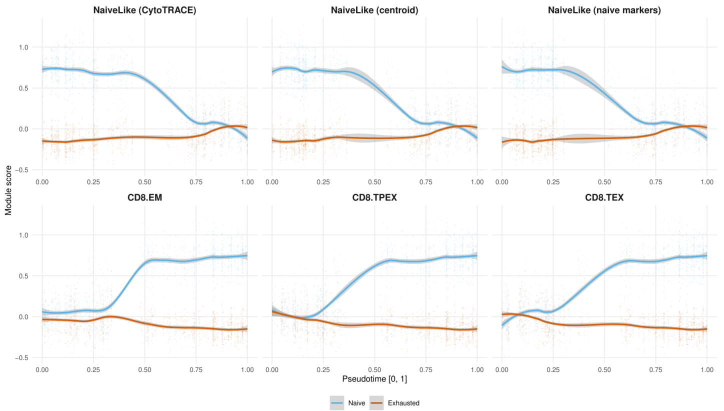

C

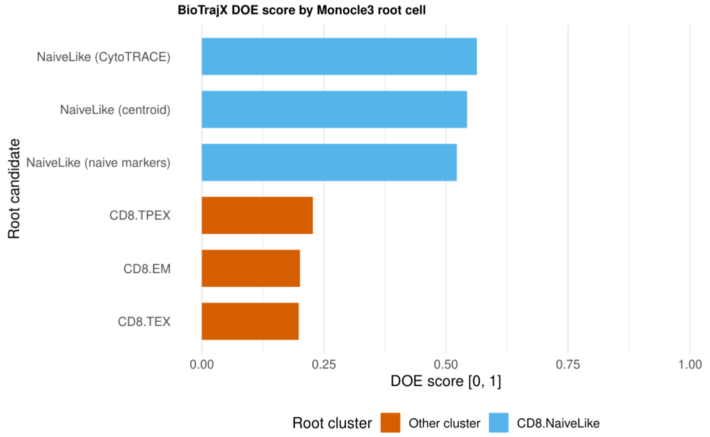

D

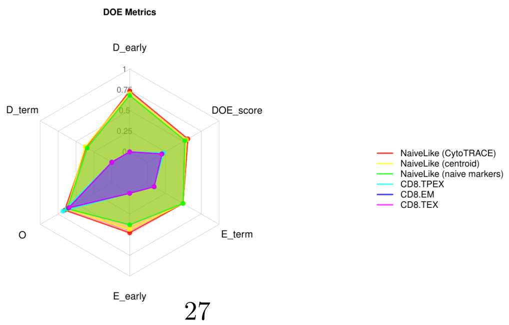
